# α-Synuclein strain dynamics predict cognitive transitions in Parkinson’s disease

**DOI:** 10.1101/2024.10.22.619694

**Authors:** Kundlik Gadhave, Ning Wang, Kyungdo Kim, Enquan Xu, Xiaodi Zhang, Hanyu Li, Jacob Deyell, Jun Yang, Anthony Wang, Youngjae Cha, Ramhari Kumbhar, Haiqing Liu, Lili Niu, Rong Chen, Shu Zhang, Catherine C. Bakker, Lingtao Jin, Yajie Liang, Mingyao Ying, Brenna Cholerton, Vadim Zipunnikov, Nikolay Bliznyuk, Joseph F. Quinn, Kathryn A. Chung, Amie L. Hiller, Kimmy Su, Shu-Ching Hu, Thomas J. Montine, Shizhong Han, Cyrus P. Zabetian, Chan Hyun Na, Valina L. Dawson, Ted M. Dawson, Liana S. Rosenthal, Xiaobo Mao

## Abstract

α-Synuclein (α-syn) strains can serve as discriminators between Parkinson’s disease (PD) and related α-synucleinopathies. The relationship between α-syn strain dynamics and clinical performance as patients transition from normal cognition (NC) to cognitive impairment (CI) is not known. Here, we show that the biophysical properties and neurotoxicity of α-syn strains change as PD cognitive status transitions from NC to mild cognitive impairment (PD-MCI) and dementia (PD-D). Both cross-sectional and longitudinal analyses reveal distinct α-syn strains in PD patients correlating to their level of cognitive impairment. Machine learning (ML) was employed to achieve high classification accuracy. The combination of thioflavin T (ThT) maximal fluorescence intensity (mfi), max slope of rise curve (forming rate), lag time (t_lag_), 20% time (t_20_), and half-time (t_50_), dynamic light scattering (DLS) (peak number, ½ peak size, ½ peak intensity) and neurotoxicity together with demographic variables for model training yielded superior performance (89∼99% accuracy in the 4- and 2- classification schema) compared to individual features alone in classifying cognitive status. For the longitudinal study, DLS peak number emerged as the strongest predictor of cognitive transition (HR = 0.12, *P* = 0.002), with the optimal predictive model combining DLS peak number, sex, education, DLS peak 1 size, and DLS peak 2 polydispersity achieving high accuracy (C-index of ∼93%). This study presents evidence that individuals with PD have different α-syn strains correlating to their cognitive status and highlights the potential of α-syn strain dynamics to guide future diagnosis, management, and stratification of PD patients.

**One Sentence Summary:** Distinct features of α-syn strains change with cognitive decline in Parkinson’s disease and AI-based analysis incorporating these combined characteristics serves as a powerful tool for PD clinical stratification.

## Introduction

Misfolded α-synuclein (α-syn) is a central pathological feature underlying the motor and cognitive changes in individuals with Parkinson’s disease (PD). Seed amplification assays (SAAs) targeting misfolded α-syn have emerged as highly sensitive and specific diagnostic tools capable of distinguishing PD from related synucleinopathies such as multiple system atrophy (MSA)^1–3^. Progression biomarkers directly related to PD-pathophysiology, particularly those predicting cognitive decline are needed to improve patient care, facilitate clinical trial cohort selection, and aid in the development of new therapeutics. Disease progression in PD includes greater motor impairment over time, often coupled with the worsening of non-motor symptoms, including cognitive change. Patients initially demonstrate limited or no cognitive changes early in the disease (PD-NC) and later develop cognitive impairment (PD-CI), including PD-MCI (mild CI) and then PD-D (dementia). The rate and severity of this cognitive change are highly variable between patients, which has been attributed, in part, to the presence of other proteinopathies at autopsy^4^ and the extent of α-syn cortical burden^5^.

Emerging data from α-syn SAA research show its expanding utility in characterizing PD cognitive stages and predicting cognitive change. Comparative studies of CSF-derived α-syn seeds in PD-NC and PD-D have identified clear differences in kinetic parameters such as lag phase and ThT fluorescence amplitude, supporting their use as stage-specific indicators^6^. Longitudinal analyses further show that α-syn aggregation alterations can be detected in individuals who later transition to PD-D, indicating predictive value at early stages^7^. Plasma-derived α-syn seeds exhibit similar stage-dependent kinetic differences, providing a minimally invasive approach to cognitive stratification^8^. These findings have been replicated across independent cohorts^9^, and recent work combining kinetic and structural readouts shows that specific α-syn strain features can signal impending cognitive decline up to one year before clinical PD-D diagnosis^10^.

We theorized that if α-syn strains underlie the heterogeneity of α-synucleinopathies, it is possible that α-syn strains change with the different cognitive stages of PD and that α-syn strain properties differ amongst individuals and within an individual over time. To investigate this possibility, we collected cerebrospinal fluid (CSF) samples from clinically well-characterized patients from two independent cohorts and characterized the biophysical, biochemical, and cellular properties of the amplified α-syn aggregates in relation to the patients’ clinical characteristics. Notably, we found that α-syn exhibits different aggregation and biophysical properties among PD patients as they transition from normal cognition (NC) to cognitive impairment (CI). These findings indicate that cross-sectional and longitudinal characterization of α-syn strain properties may be of value in the management of PD patients as they transition from NC to CI. Together, these results move the field from binary SAA diagnostics toward quantitative, prognostic strain readouts for PD cognitive decline, with a defined ∼1-year prediction window for PD-NC to PD-MCI transition.

## Results

### α-Synuclein strains change in relation to cognitive status

Protein misfolding cyclic amplification (PMCA) was used as the SAA from cerebrospinal fluid (CSF) obtained in a cross-sectional manner from PD patients and controls from Johns Hopkins (JHU, Cohort I) and the Pacific Udall Center (PUC, Cohort II) (**Fig. 1a**). A series of biophysical and cellular studies including Thioflavin T (ThT) assessments (**Fig. 1a**) were performed to characterize the aggregated α-syn features across diagnostic groups (**Extended Data Tables 1-3**). Multivariable logistic regression, adjusting for sex, age, education, and disease duration, revealed that multiple α-syn strain properties significantly predicted cognitive status (PD-NC vs PD-CI, including PD-MCI and PD-D), with cognitive group determined by consensus conference diagnosis (**Supplementary Tables S24–S26**).

**Figure 1.**
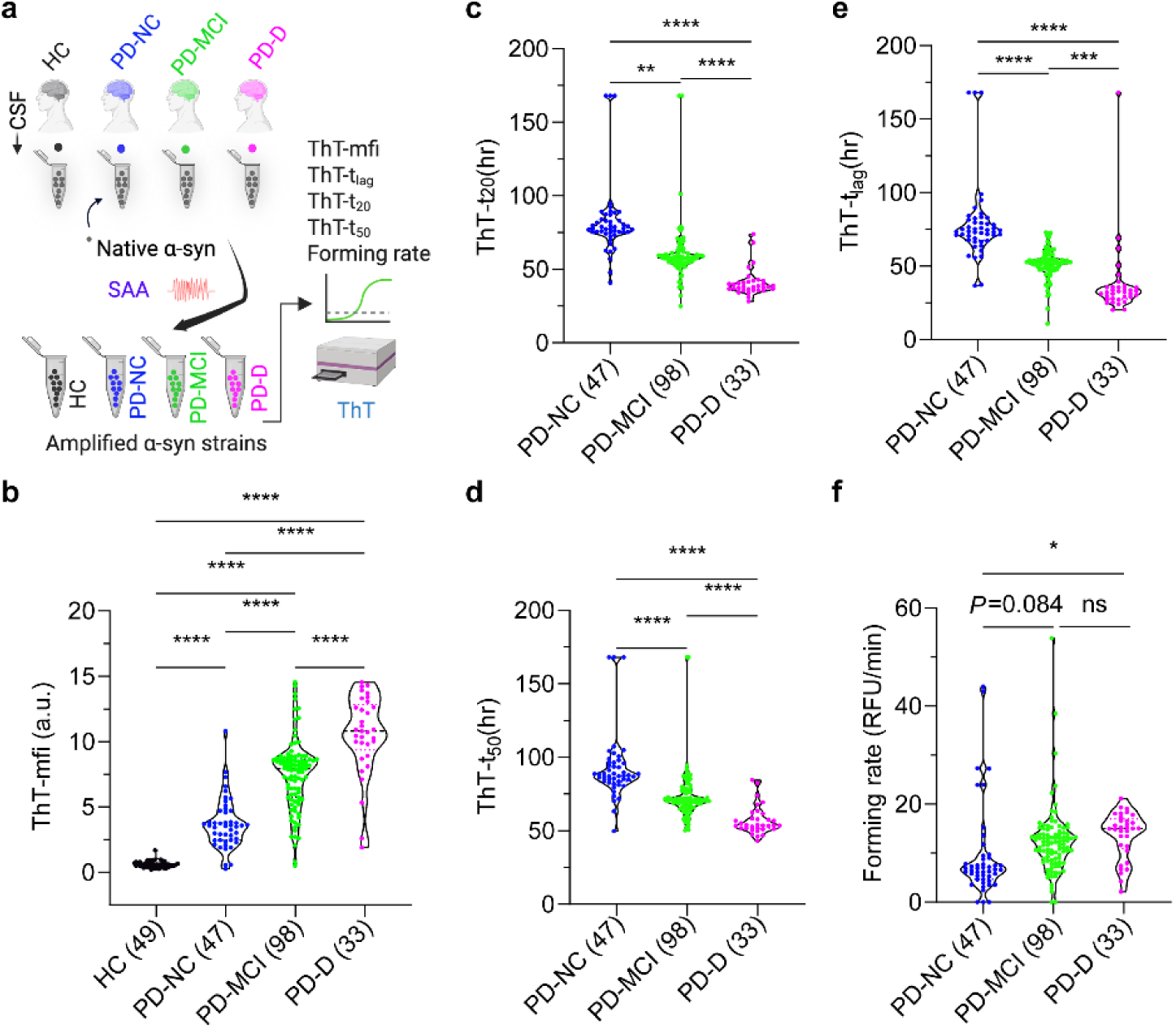
Differentiating amplified α-syn strains via Thioflavin T (ThT) derived from patients with PD-NC, PD-MCI, and PD-D. (**a**) Schematic representation of α-syn amplification and characterization using ThT assay. In these groups: HC (health control), PD-NC (normal cognition), PD-MCI (mild cognitive impairment), PD-D (dementia), and CSF samples were amplified with SAA and the ThT assay was performed. (**b**) ThT-mfi (maximal fluorescence intensity) of CSF-SAA samples. HC (*n* = 49), PD-NC (*n* = 47), PD-MCI (*n* = 98), and PD-D (*n* = 33). (**c**) ThT-t_20_ (time reach to 20% ThT-mfi of each sample) of CSF-SAA samples. PD-NC (*n* = 47), PD-MCI (*n* = 98), and PD-D (*n* = 33). (**d**) ThT-t_50_ (time at which aggregation is 50% complete) of CSF-SAA samples. PD-NC (*n* = 47), PD-MCI (*n* = 98), and PD-D (*n* = 33). (**e**) ThT-t_lag_ (time at which aggregation started) of CSF-SAA samples. PD-NC (*n* = 47), PD-MCI (*n* = 98), and PD-D (*n* = 33). (**f**) Forming rate of CSF-SAA samples. PD-NC (*n* = 47), PD-MCI (*n* = 98), and PD-D (*n* = 33). Data are mean ± SEM. The statistical significance was evaluated via one-way ANOVA with Tukey’s multiple comparisons test. No significant difference (ns) *P* > 0.05, \*\**P* < 0.01, \*\*\**P* < 0.001, \*\*\*\**P* < 0.0001. Every dot indicates an individual biological sample measured in duplicate.

### ThT features distinguish α-synuclein strains among the cognitive groups

The ThT results showed that both the maximal fluorescence intensity (mfi), Lag time (t_lag_), max slope of ThT rise curve (forming rate) and time at which aggregation is 20% complete (t_20_) and 50% complete (t_50_) significantly predict the odds of a PD-CI diagnosis over PD-NC when controlling for sex, education, age at baseline visit, and disease duration (**Supplementary Tables 24-26**). ThT-mfi was significantly higher in PD-D, then decreased in a step-wise fashion with significant differences between PD-D and PD-MCI, PD-MCI and PD-NC, and PD-NC and HC (**Fig. 1b**). The ThT t_20_ differed among the cognitive groups. The PD-D group spent the least ThT-t_20_ compared to the PD-MCI and PD-NC groups (**Fig. 1c**). The PD-MCI group spent less ThT-t_lag_ than the PD-NC group (**Fig. 1c**). There is no ThT-t_20_ for the HC group due to the minimal fluorescence intensity (**Fig. 1b,c**). The ThT-t_50_ and ThT-t_lag_ of PD-D was significantly lower than ThT-t_50_ and ThT-t_lag_ of the PD-MCI and PD-NC groups (**Fig. 1d,e**). Forming rate, an emerging quantitative metric for assessing pathological protein seeding activity in the serum, has shown promise as a liquid biomarker for Lewy body diseases^2^. The forming rate of PD-D was significantly higher than the forming rate of PD-NC group but not PD-MCI group (**Fig. 1f**). There is an elevated trend of forming rate in the PD-MCI group compared to the PD-NC group (*P* = 0.08) (**Fig. 1f**). Similar results for PD, PD-MCI and PD-D were observed when the analyses were separated into cohort I and cohort II (**Supplementary Tables S24–S26**).

### Dynamic light scattering (DLS) properties of α-synuclein strains

DLS was utilized to assess the size profiles (i.e., peak number, intensity, size) of amplified α-syn strains (**Fig. 2a**). The typical DLS spectra of both the PD-MCI and PD-D groups showed significantly greater homogeneity with one peak compared to the PD-NC and HC groups, which exhibited two peaks (**Fig. 2b,c, and Extended Data figure 2e,i**). Peak size was not significantly different between PDD and PD-MCI, but decreased in a step-wise fashion with significant differences between PD-D or PD-MCI versus PD-NC and HC (**Fig. 2d, e**). Peak intensity was also not different between PD-D and PD-MCI, but increased in a step-wise fashion with significant differences between PD-D or PD-MCI versus PD-NC and HC (**Fig. 2f,g**), which is similar to DLS percent polydispersity from polydispersity index (pd%) of peak 1 and peak 2 (**Fig. 2h,i**). Differences in the peak size and peak intensity were also observed when the analyses were separated into cohort I and cohort II (**Extended Data figure 2**). Evaluations of these architectural features demonstrated that the peak number, particle dimensions, and intensity distributions provide critical diagnostic weight that significantly increases the probability of identifying PD-CI over PD-NC, even when controlling for motor symptoms and demographic factors (**Supplementary Tables 24-26**).

**Figure 2.**
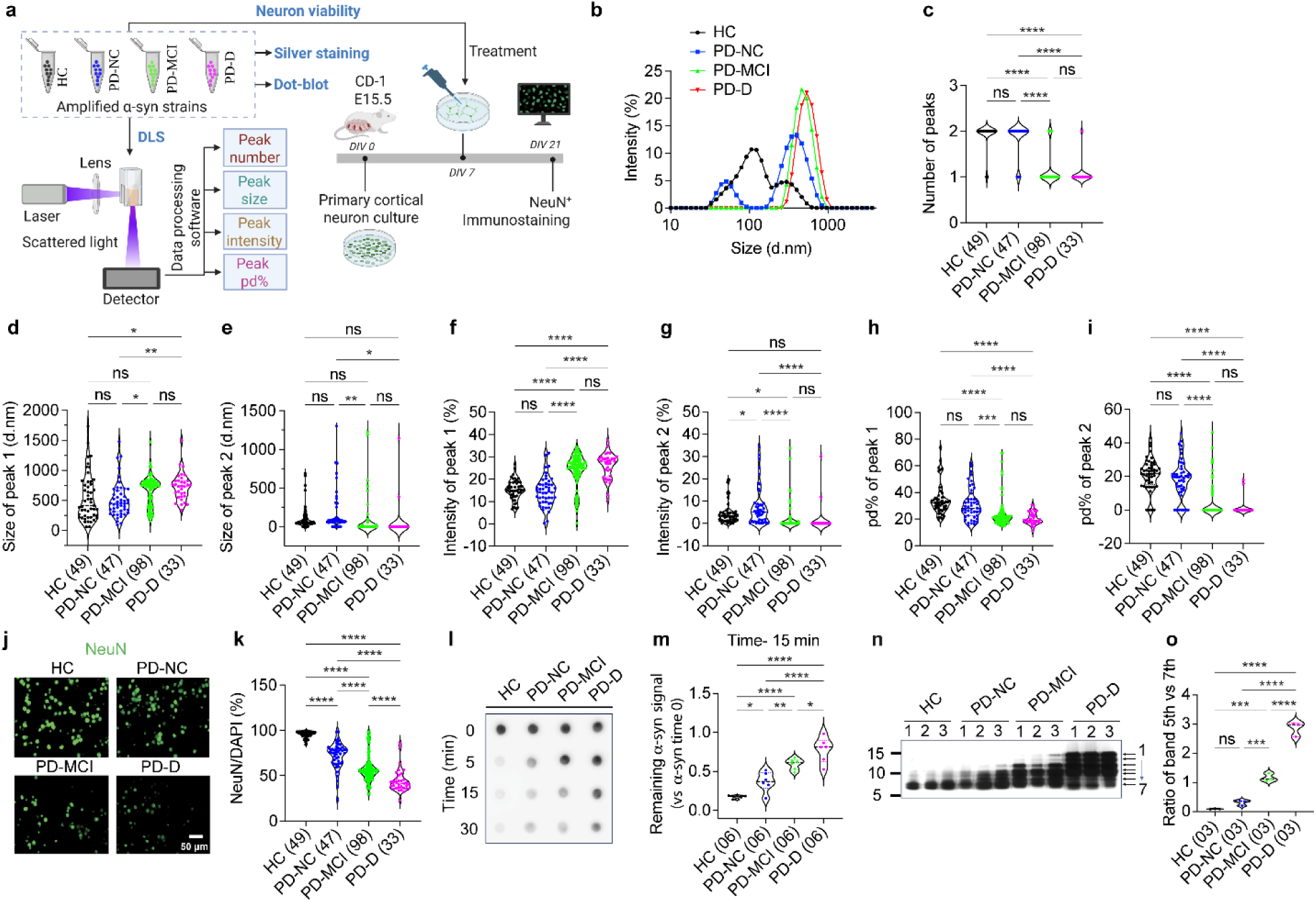
Differentiating α-syn strains using dynamic light scattering (DLS), cell-based, and biochemical assays. (**a**) Schematic representation of the characterization of α-syn strains from HC, PD-NC, PD-MCI, and PD-D. DLS data processing provides peak number, peak size, peak intensities and peak pd%. Neuronal culture were used to assess neurotoxicity. (**b**) DLS spectra of amplified α-syn strains from HC, PD-NC, PD-MCI, and PD-D. (**c**) The number of DLS for HC (*n* = 49), PD-NC (*n* = 47), PD-MCI (*n* = 98) and PD-D (*n* = 33). (**d**) Size of peak 1 from DLS for amplified α-syn strains from HC, PD-NC, PD-MCI, and PD-D. (**e**) Size of peak 2 from DLS for amplified α-syn strains from HC, PD-NC, PD-MCI, and PD-D. (**f**) Intensity of peak 1 from DLS for amplified α-syn strains from HC, PD-NC, PD-MCI, and PD-D. (**g**) Intensity of peak 2 from DLS for amplified α-syn strains from HC, PD-NC, PD-MCI, and PD-D. (**h**) pd% of peak 1 from DLS for amplified α-syn strains from HC, PD-NC, PD-MCI, and PD-D. (**i**) pd% of peak 2 from DLS for amplified α-syn strains from HC, PD-NC, PD-MCI, and PD-D. (**j & k**) Neurotoxicity of α-syn strains assessed with the immunostaining and quantification of anti-NeuN (neuronal nuclei marker). HC (*n* = 49), PD-NC (*n* = 47), PD-MCI (*n* = 98) and PD-D (*n* = 33). Scale bar, 50 µm. (**l**) α-syn dot-blot immunostaining after proteinase K (PK)-digestion and **(m)** quantification. HC (*n* = 6), PD-NC (*n* = 6), PD-MCI (*n* = 6) and PD-D (*n* = 6). (**n**) SDS–PAGE followed by silver staining of PK-digested α-syn strains and (**o**) quantification. HC (*n* = 3), PD-NC (*n* = 3), PD-MCI (*n* = 3) and PD-D (*n* = 3). (**c-i, k, m, o**) Each dot represents an individual biological sample. The violin plot shows all the points. Data are presented as the mean ± SEM. The statistical significance was evaluated via one-way ANOVA with Tukey’s multiple comparisons test. No significant difference (ns) *P* > 0.05, \**P* < 0.05, \*\**P* < 0.01, \*\*\**P* < 0.001, \*\*\*\**P* < 0.0001.

### α-Synuclein strains are differentially neurotoxic

The amplified ɑ-syn strains were applied to primary mouse cortical neurons and neurotoxicity was assessed by quantifying NeuN immunostaining (**Fig. 2j**). Consistent with the delineation of an α-syn strain exhibiting markedly heightened neurotoxic potency^11^, the amplified ɑ-syn strains from the PD-NC group exhibited more neurotoxicity than the HC group (**Fig. 2j,k**). The PD-MCI group showed significantly increased neurotoxicity compared to the PD-NC group, and the PD-D group exhibited the highest neurotoxicity (**Fig. 2k**). Similar results for HC, PD-NC, PD-MCI and PD-D were observed when the analyses were separated into cohort I and cohort II (**Supplementary Table 24**). Quantitative analysis established that the neurotoxic feature significantly correlates with cognitive decline, showing that increased toxicity levels independently heighten the odds of a clinical diagnosis of PD-CI (including PD-MCI and PD-D) over PD-NC after adjusting for demographic and clinical variables (**Supplementary Tables 24-26**).

### α-Synuclein strains exhibit different Proteinase K (PK) digestion patterns

PK digestion was performed on the amplified ɑ-syn strains over time, and followed with a dot-blot assay for ɑ-syn immunoreactivity (**Fig. 2l,m**) and silver staining (**Fig. 2n,o**) as previously described^12,13^. In the dot-blot assay, the PD-D strain showed the strongest resistance to PK digestion as evidenced by the remaining ɑ-syn signal (**Fig. 2l, m and Extended Data figure 2u, v**); the PD-MCI strain exhibited less resistance than PD-D strain, but more resistance than PD-NC strain (**Fig. 2l, m and Extended Data figure 2u,v**); the PD-NC strain exhibited mild resistance to PK digestion; the HC strain has minimal resistance to PK digestion (**Fig. 2l, m and Extended Data figure 2u,v**). To assess the digested band patterns in response to PK digestion via silver staining, the digested bands between the 5^th^ and 7^th^ bands were quantified (**Fig. 2n**). The results showed significant differences between the PD-D and PD-MCI strains (**Fig. 2n,o**), and between the PD-MCI and PD-NC strains. However, there was no difference between the PD-NC and HC strains (**Fig. 2o**). Similar results for HC, PD-NC, PD-MCI, and PD-D were observed when the analyses were separated into cohort I and cohort II (**Extended Data figure 2**). The structural resilience of these strains, particularly the amount of remaining ɑ-syn signal after 15 minutes of digestion, serves as a biochemical indicator of conformational stability and protease resistance^14^.

### Predicting PD clinical phenotypes using machine learning analysis of amplified ɑ-synuclein strains

We developed ML classifiers to distinguish HC, PD-NC, PD-MCI, and PD-D using 13 biochemical features (5 ThT, 7 DLS, and 1 Neurotoxicity features) together with demographic variables (age, sex, education, and disease duration). Seven classification algorithms (classifiers) were evaluated: CatBoost (CB), Gradient Boosting Decision Trees (GBDT), XGBoost, Random Forest, Decision Tree, Extra Trees (ET), and Logistic Regression with L1 regularization. For the 4-class classification task, CB was excluded due to compatibility issues with multi-class settings, resulting in six classifiers for that task. For feature evaluation we tested predefined sets (each of the 13 individual features, the 5 ThT-feature set, the 7 DLS-feature set, and the full 13-feature set) and also applied a Top-K approach: Random Forest feature importance was used to rank features and create Top-K subsets (k = 1, 2, 3, 5, 7, 10, 13, 15, and all features) for targeted evaluation. For each dataset and feature combination, we performed cross-validated comparisons across all seven classifiers to identify the best-performing algorithm, followed by feature importance analysis to assess the contribution of individual features.

For the ML classification, dataset 0 comprised 227 individuals from the Cohort I and Cohort II (**Fig. 3a**). Each case was defined by 13 distinct features and up to 4 demographic variables and was assigned a label that indicates their disease/cognitive group, categorized as HC (49 cases), PD-NC (47 cases), PD-MCI (98 cases), or PD-D (33 cases). For the HC vs PD classification (Task 1), 16 features were used (13 biomarkers + age, sex, and education), with disease duration excluded because all HC subjects have a disease duration of 0 by definition, making it a trivially discriminating variable. For the PD-NC vs PD-CI classification (Task 2) and the 4-class classification (Task 3), all 17 features (13 biomarkers + age, sex, education, and disease duration) were included, as disease duration provides meaningful variation within PD patient subgroups. From dataset 0, two additional datasets were created for binary classification: the first dataset (**dataset 1**) consolidated the PD-NC, PD-MCI, and PD-D groups into a single PD group (178 cases) while keeping the HC group (49 cases) the same (**Fig. 3a**). The second dataset (**dataset 2**) excluded all HC records and aggregated PD-MCI and PD-D patients into a single CI category, resulting in NC (47 cases) vs CI (131 cases) groups for classification (**Fig. 3a**).

**Figure 3.**
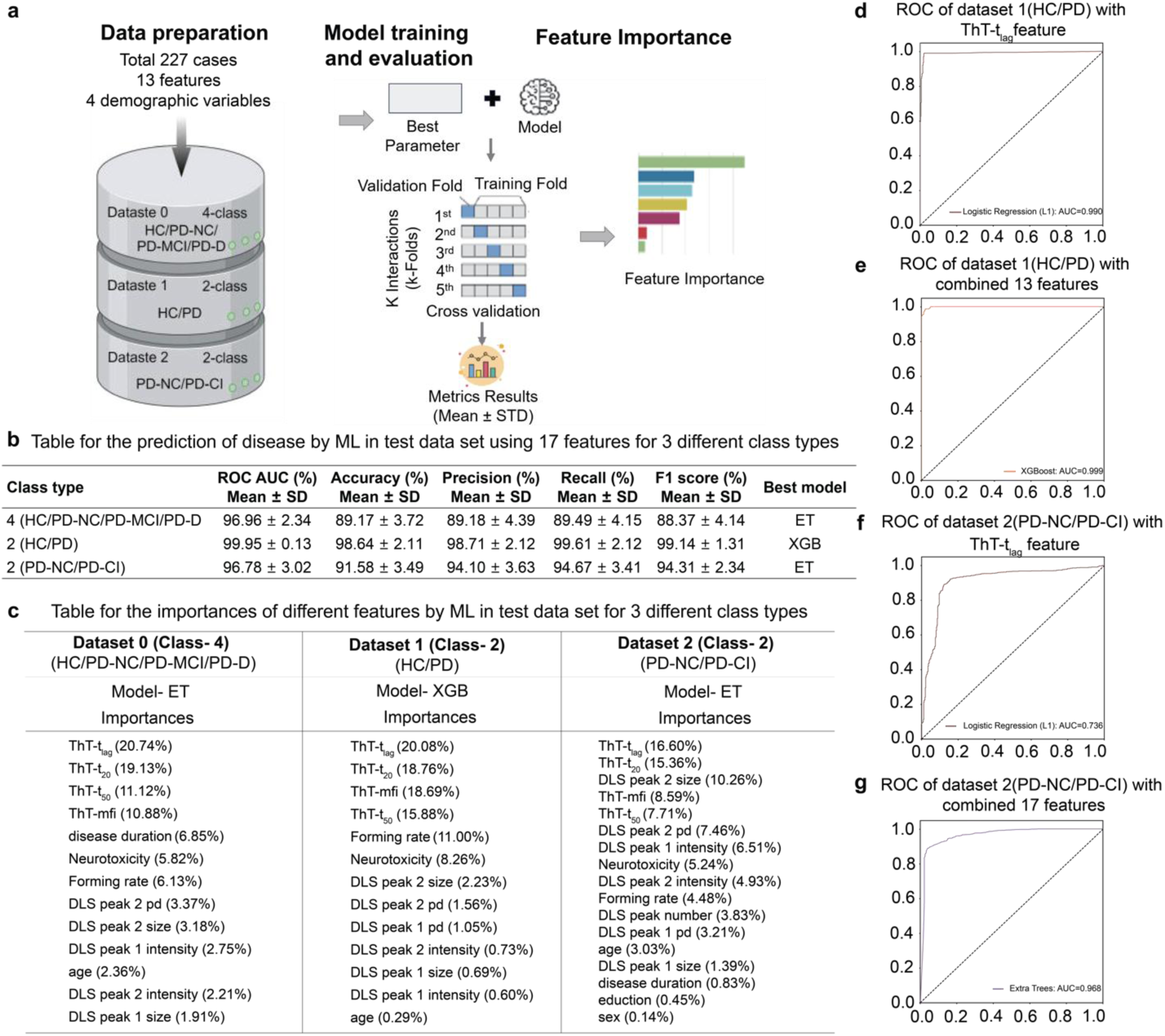
Machine Learning (ML) predicted disease using ThT, DLS, and neurotoxicity data. (**a**) Schematic representation of how ML was used to predict disease using ThT, DLS, neurotoxicity, and demographic. (**b**) Table for the prediction of disease by ML using the best model for 3 different class types. (**c**) Table for the importances of different features by ML for 3 different class types. (**d**) ROC for dataset 1 (HC vs PD) with the ThT-t_lag_ feature using the Logistic Regression (L1) model. (**e**) ROC for dataset 1 (HC vs PD) with combined 13 features using the XGBoost model. (**f**) ROC for the dataset 2 (PD-NC vs PD-CI) with ThT-t_lag_ feature using Logistic Regression (L1) model. (**g**) ROC for dataset 2 (PD-NC vs PD-CI) with combined 17 features using the ET model.

We applied a Repeated Stratified K-fold cross-validation procedure to train and evaluate model performance. By using a fixed random state, we ensured that the data splits were identical for each model. Scoring metrics included accuracy, precision, recall, F1 score, and Area Under the Receiver Operating Characteristic Curve (AUC) (**Fig. 3d-g, Extended Data figures 3 and 4**). For multi-class classification, we calculated AUC using ‘macro’ averaging and a ‘one-vs-rest’ strategy. (**Fig. 3a**). Post-evaluation, feature importance was extracted from the models and averaged across all classifiers (**Fig. 3b,c**). To assess generalizability, Leave-One-Cohort-Out (LOCO) validation was performed using the best-performing model for each task (**Supplementary Table 15**).

### Superiority of 13-features and the importance of ThT in 4-class classification of HC vs. PD-NC vs. PD-MCI vs. PD-D

Following the two complementary selection strategies (predefined sets and RF-guided Top-K) as described in Supplementary Methods, we prepared various combinations of feature sets, followed by conducting multivariate analysis of the feature sets using the algorithms described above. The Extra Trees model trained on the top 13 RF-ranked features emerged as the best performer for 4-class discrimination (see below and **Supplementary Table 10**); we therefore focus on this configuration while reporting other comparisons for context. The ET model, which utilized the full set of 13 features (ThT-t_lag_, ThT-t_20_, ThT-t_50_, ThT-mfi, disease duration, forming rate, Neurotoxicity, DLS-peak-2-pd%, DLS-peak-2-size, DLS-peak-1-intensity, age, DLS-peak-2-intensity, DLS-peak-1-size), achieved the highest mean AUC of 96.96% ± 2.34% (**Fig. 3b** and **Supplementary Table 10**). This model also demonstrated high accuracy (89.17% ± 3.72%), precision (89.18% ± 4.39%), recall (88.49% ± 4.15%), and F1 score (88.37% ± 4.14%) (**Fig. 3b** and **Supplementary Table 10**), indicating its reliability and effectiveness in classifying the 4 classes (HC, PD-NC, PD-MCI, and PD-D). The aggregated confusion matrix across all cross-validation folds confirmed robust discrimination, with HC and PD-D cases showing particularly high classification rates (**Supplementary Table 13 and 14**).

The 5 ThT-feature set also performed well across all models, but not as well as the 13-feature set (**Supplementary Table 12)**. The Extra Trees model with the 5 ThT-feature set achieved a mean AUC of 94.90% ± 2.15%, accuracy of 83.35% ± 4.19%, precision of 83.70 ± 5.16%, recall of 83.05% ± 4.75%, and F1 score of 82.56% ± 4.67% (**Supplementary Table 12**). This suggests that the inclusion of five extra features can enhance the classification performance.

In terms of individual features, the five ThT-related features (ThT-mfi, ThT-t_20_, ThT-t_50_, ThT-t_lag_, forming rate) demonstrate high performance across all models, with a mean AUC between 68 to 95% (**Supplementary Table 9**). This suggests that ThT-related features are important indicators for the 4-class classification. Among these, the ThT-t_lag_ feature consistently showed the highest performance across all models, with a mean AUC ranging from 81.64% to 94.72%, indicating that the lag time of ThT fluorescence is a particularly significant factor (**Supplementary Table 9**).

The seven DLS-related features showed more varied performance. The DLS-peak-1-size resulted in notably lower performance, with mean AUC consistently below 64% across all models, ranging from 50.57% to 63.53% (**Supplementary Table 9**). Conversely, the DLS-peak-2-size feature showed relatively high performance, with mean AUC mostly around 65-75% (**Supplementary Table 9**). This indicates that the size of the second DLS peak could be a more important factor in the classification than the first peak.

The Neurotoxicity feature also achieved a relatively high mean AUC of 87.31% ± 3.65% with the logistic regression model, while other models’ mean AUC were all above 70% (**Supplementary Table 9**), indicating that neurotoxicity could also be a significant factor in classifying the cognitive groups.

The feature importance analysis revealed that the top four features that consistently showed significance across all models were ThT-t_lag_, ThT-t_20_, ThT-t_50,_ and ThT-mfi (**Fig. 3c**). ThT-t_lag_ was consistently ranked highest across both the 5 ThT-feature set and 13-feature set in multiple models, with the highest importance score of 20.74% in the ET model (**Fig. 3c, Supplementary Table 11**). ThT- t_20_ also emerged as an essential feature across all models, with the importance given by the ET model for the 13-feature set of 19.13% (**Fig. 3c**). ThT-t_50_ was another feature that consistently showed substantial importance across all models, with the importance score of 11.12% in the ET model for the 13-feature set (**Fig. 3c**). Beyond these 3 features, other features including ThT-mfi (10.88% in ET model), disease duration (6.85% in ET model), forming rate (6.13% in ET model), neurotoxicity (5.82% in ET model), and those related to the DLS peak also provided valuable insights, particularly when considering a more diverse set of factors in the 13-feature set.

### Superiority of the 13-features and the importance of the ThT features in binary classification of HC vs. PD

Using the two complementary feature selection strategies described in the Supplementary Methods, we prioritized the RF-ranked top 13 features for the HC vs PD binary analyses and report those results alongside ThT-only comparisons for context. The binary classification task to distinguish between HC and PD patients yielded notable results. The performance of models, however, varied considerably depending on the features used. When using individual features, most models using ThT-related features achieved an AUC above 90%, with ThT-t_lag_ reaching up to 99.30% (Logistic Regression) (**Supplementary Table 1**). Conversely, models using the ‘DLS-peak-1-intensity’ only achieved an AUC ranging from 66.38% to 81.92% (**Supplementary Table 1**). This underscores the impact of feature selection on performance and highlights the discriminative power of ThT features in distinguishing between HC and PD.

When the 5 ThT-feature set was used, the models achieved comparable AUC relative to those using individual ThT features. The ROC AUC of these models ranged from 95.45% to 99.61%, indicating a strong performance in terms of both sensitivity and specificity (**Supplementary Table 4**). However, the multi-feature set models consistently outperformed the others. The highest mean AUC of 99.95% ± 0.13% was achieved by the XGBoost model using the top 13 features (**Supplementary Table 2**). Similarly, the accuracy, precision, recall, and F1 score were also highest for the 10-feature set, with the Extra Trees model (top-7 features) achieving a mean accuracy of 98.94% ± 1.40%, precision of 99.45% ± 1.11%, recall of 99.21% ± 1.38%, and F1 score of 99.32% ± 0.90% (**Fig. 3b**).

Feature importance analysis indicates that ThT-t_lag_, ThT-t_20,_ ThT-mfi, and ThT-t_50_ are the most important features in the 13-feature set (**Fig. 3c, Supplementary Table 3**). The top six features by importance were ThT-t_lag_ (20.08%), ThT-t_20_ (18.76%), ThT-mfi (18.69%), ThT-t_50_ (15.88%), forming rate (11.00%), and neurotoxicity (8.26%). This suggests that a combination of these 10 features provides a more accurate and reliable classification of PD and HC than any individual feature or the 5 ThT-feature set.

The 5ThT-feature set also performed better than most of the other feature sets, except for the 10-feature set. This indicates that while combining 5 ThT features together was effective, the inclusion of additional features in the 10-feature set further improves the model’s performance.

### Superiority of 17-features and the importance of ThT features in binary classification of NC vs. CI (Dataset 2)

Applying the same evaluation framework to the NC vs. CI classification task, we assessed both individual features and comprehensive feature combinations. The binary classification task to distinguish between NC and CI of PD also yielded notable results. Upon evaluating the performance of models utilizing individual features, the ‘ThT-t_lag_’ feature, when incorporated with the Logistic Regression model, exhibited the highest mean AUC of 92.19% ±5.29% (**Supplementary Table 5**). This was closely followed by the ‘ThT-t_20_’ feature, achieving a mean AUC of 91.90% ± 5.46% with the Logisitic Regression model (**Supplementary Table 5**). The ‘DLS peak 2 size’ feature also demonstrated a robust predictive capability, with the CatBoost model achieving a mean AUC of 87.60% ± 6.62% and a consistent mean accuracy of 87.03% ± 4.83% (**Supplementary Table 5**).

The 5 ThT-feature set, when used with the CB model, achieved a mean AUC of 95.54% ± 3.15% and a mean accuracy of 88.60% ± 5.21% (**Supplementary Table 8**), which is better than the best individual feature model. However, a significant improvement was observed when the 17-feature set was employed. The Extra Trees model, in particular, achieved the highest mean AUC, reaching 96.78% ± 3.02% and a mean accuracy of 91.58% ± 3.49% (**Fig. 3b** and **Supplementary Table 6**). The feature importance analysis revealed that ThT-t_lag_, ThT-t_20_, and DLS peak 2 size consistently ranked high in terms of importance across different models (**Supplementary Table 7**). Furthermore, the DLS-peak-number achieved a mean AUC of 86.44% ± 6.47% and accuracy of 87.08% ± 5.27% (**Supplementary Table 5**). This indicates that while the DLS-peak-number was a strong predictor on its own, its predictive power can be enhanced when it is used in conjunction with other features. This indicates that the 17-feature set provided the most accurate predictions. Notably, the DLS peak 2 size feature ranked third in overall feature importance (10.26%), demonstrating that DLS-derived architectural features provide complementary discriminative information beyond ThT kinetics for the PD-NC vs. PD-CI classification.

These analyses taken together highlight the potential of ML classifiers in differentiating various PD stages. For the individual features, the ThT-related features consistently showed high performance across all models, with ThT-t_lag_ being the most significant. This suggests that ThT features are important indicators for distinguishing among different levels of cognitive impairment in PD. The DLS-related features and the neurotoxicity feature provided valuable insights, with the DLS-peak-2-size and DLS-peak-number demonstrating particularly strong predictive capabilities in the binary classification of NC vs. CI.

The use of expanded feature sets incorporating both biomarker and demographic variables leads to the highest accuracy in all 3 classification tasks, indicating that a more comprehensive set of informative features improves classification performance. Leave-One-Cohort-Out validation further confirmed generalizability: HC vs PD-D classification maintained > 99% AUC in both cross-cohort directions, while 4-class classification achieved AUC of 93.16% (Cohort I to Cohort II) and 99.51% (Cohort II to Cohort I) (**Supplementary Table 15**).

### α-Synuclein strains change in relation to cognitive status in the same individual

To determine whether α-syn strains change within the same individuals, the CSF samples from the same individuals obtained at the initial and last visits (either a 3-, 4-, or 5-year follow-up time) of Cohort I were used to amplify α-syn strains followed by characterization and correlation studies (**Fig. 4 and 5**). ThT-mfi did not change if the cognitive status remained the same between the initial and last visits (**Fig. 4a**). That is, patients who remained HC, PD-NC, PD-MCI, or PD-D between their initial and last visits had a stable ThT-mfi that matched their cognitive strata.

**Figure 4.**
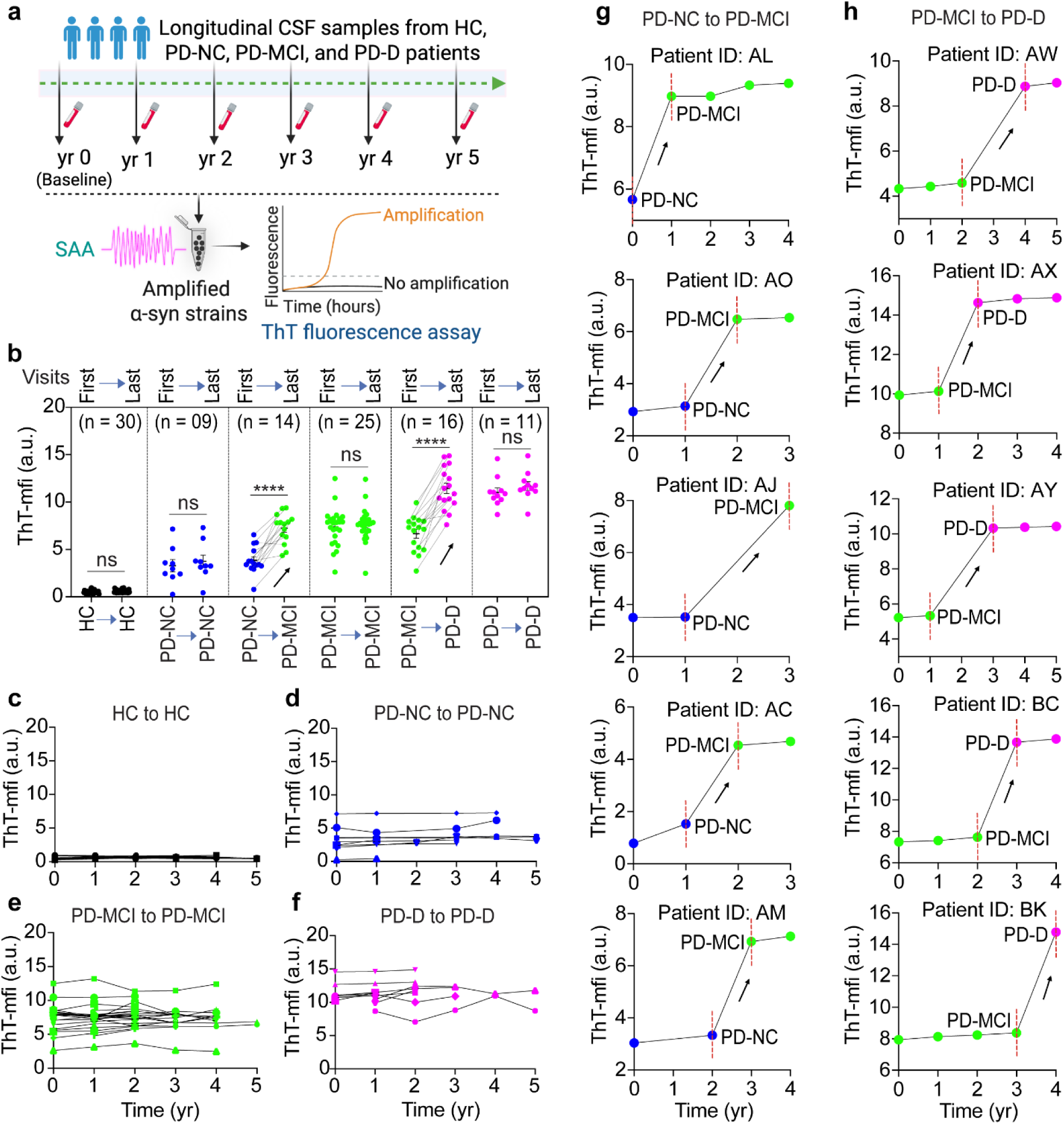
Longitudinal analysis of α-syn strains using ThT-mfi. (**a**) ThT-mfi of amplified α-syn strains from HC, PD-NC, PD-MCI, and PD-D groups between the first- and last-visit of Cohort I. (**b,c,f**) Yearly mapping of ThT-mfi of amplified α-syn strains from individuals with stable cognitive status. (**d,e**) Yearly mapping of ThT-mfi of amplified α-syn strains from individuals with changed cognitive status. The statistical significance was evaluated via one-way ANOVA with Tukey’s multiple comparisons test. No significant difference (ns) *P* > 0.05, \*\*\*\**P* < 0.0001.

**Figure 5.**
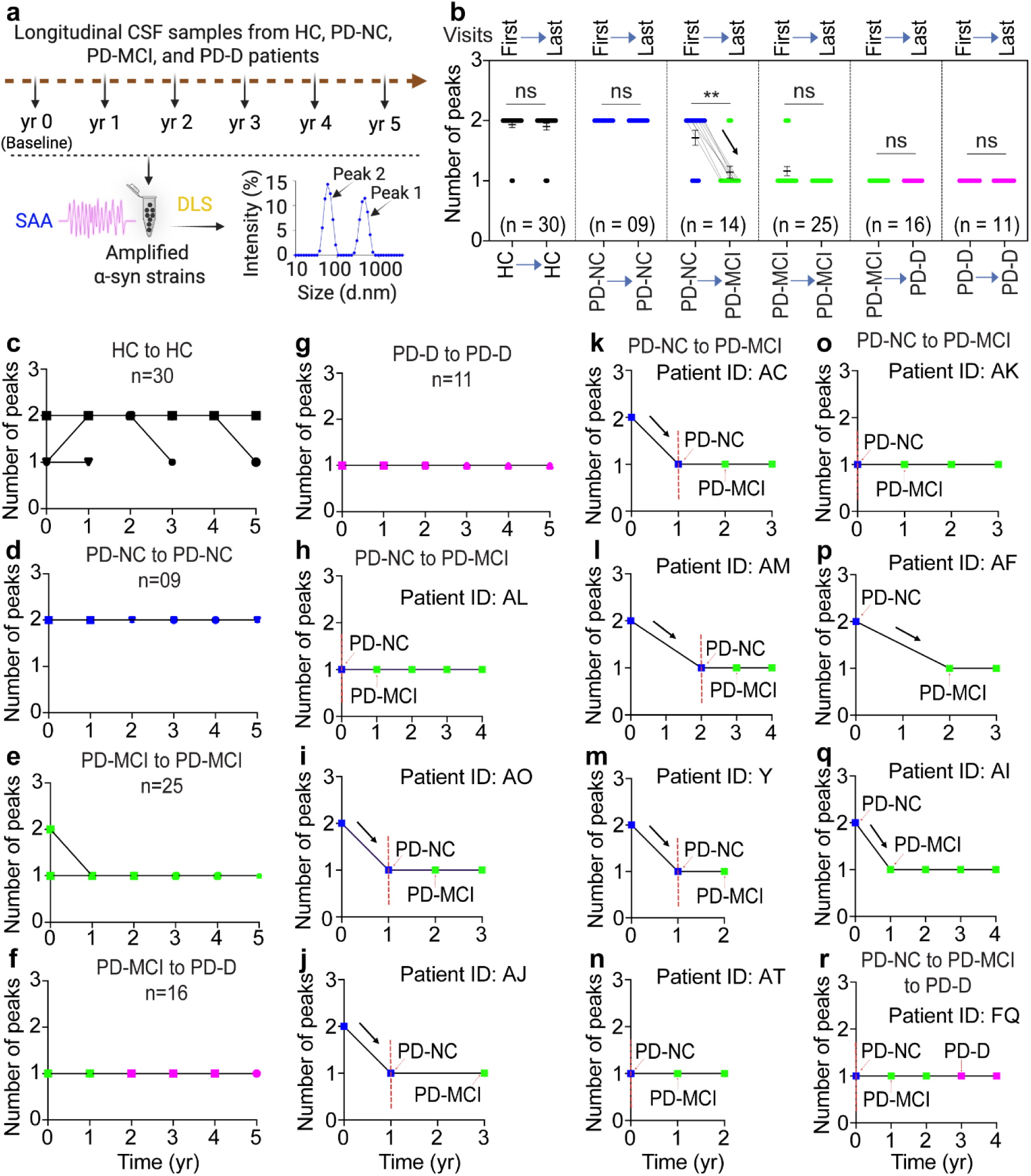
Longitudinal analysis of α-syn strains using DLS. (**a**) Schematic representation of DLS analysis of amplified α-syn strains from HC, PD-NC, PD-MCI, and PD-D groups. (**b**) Nasted plot for the peak number between the first- and last-visit in Cohort I. Different colors indicate the different stages of cognitive status. **(c,d,f,g)** Yearly mapping of the DLS peak number of amplified α-syn strains from individuals with stable cognitive status. Black: HC; blue: PD-NC; green: PD-MCI; purple: PD-D. **(e)** Yearly mapping of the DLS peak number of amplified α-syn strains from individuals with changed cognitive status.

### ThT parameters change with cognitive decline

In contrast, the ThT-mfi significantly increased in the individuals whose cognition declined by the last visit compared to the initial visit: PD-NC→PD-MCI, and PD-MCI→PD-D (**Fig. 4a**). Because the ThT results are correlated with the longitudinal cognitive decline, the time resolution was extended to determine whether the ThT profile could predict cognitive decline. In the ThT studies of α-syn strains amplified from these yearly collected longitudinal CSF samples, the results showed the ThT-mfi remained the same in those individuals without cognitive change (**Fig. 4b-f**), consistent with the ThT results of the first-last visit (**Fig. 4a**). In those individuals whose cognitive status changed, including PD-NC→PD-MCI, and PD-MCI→PD-D, the results showed that the ThT-mfi increased when cognition progressed from PD-NC to PD-MCI, and PD-MCI to PD-D (**Fig. 4b,g,h**). Similarly, ThT-t_lag_ (**Extended Data figure 6a-d and 7**) and ThT-t_50_ (**Extended Data figure 8a,c,d**) were significantly reduced in individuals whose cognition changed, PD-NC→PD-MCI, and PD-MCI→PD-D. The groups without cognitive change PD-NC→PD-NC, PD-MCI→PD-MCI and PD-D→PD-D did not show any change in ThT-t_lag_ (**Extended Data figure 6a,b,e**) or ThT-t_50_ (**Extended Data figure 8a,b,e**). These two ThT features only changed once cognition declined; it did not predate the cognitive decline.

### DLS peak number changes one year before the diagnosis of PD-MCI

DLS was used to assess longitudinal cognitive status (**Fig. 5a**). Only the group of PD-NC→PD-MCI showed a peak number change (from 2 to 1) (**Fig. 5b**). The peak number of the other groups exhibited no change, including HC→HC, PD-NC→PD-NC, PD-MCI→PD-MCI, PD-MCI→PD-D, and PD-D→PD-D (**Fig. 5b**). In the yearly mapping, the peak number remained the same in those individuals without cognitive change (**Fig. 5c,d,f,g** and **Extended Data figure 9**). In Cohort I, there were a total of 14 patients who transitioned from PD-NC to PD-MCI. Patient AG and IC exhibited an unusual fluctuating cognitive trajectory (PD-NC → PD-MCI → PD-NC) while maintaining a consistently stable DLS peak number of 2. For patient AF, the CSF sample one-year before cognitive status change to PD-MCI (year 1, PD-NC) is not available; therefore, this patient’s one-year prediction before PD-MCI is not conclusive. Among the remaining 11 patients, a striking predictive trend emerged: 09 individuals with a DLS peak number of 1 during the PD-NC stage progressed to PD-MCI within the following year, with only patients AI and AV deviating from this pattern. This strong association underscores the potential of a DLS peak number of 1 in PD-NC stage as a promising biomarker for predicting the transition from PD-NC to PD-MCI (**Fig. 5h-r** and **Extended Data figure 9**). The peak number did not change with disease duration or during the transition from PD-MCI to PD-D (**Fig. 5b,f,g** and **Extended Data figure 9**).

### Predicting Cognitive Impairment Transition in Parkinson’s Disease Using Survival Analysis with Time-Varying Covarites

Our cross-sectional analysis revealed that α-syn strain characteristics distinguish between PD-CI and PD-NC, prompting investigation of their predictive capability for PD-NC to PD-MCI transition. For this analysis, we focused on a subset of 21 patients from the Cohort I who had longitudinal CSF samples and met the criteria of PD-NC at baseline with follow-up cognitive assessments enabling evaluation of PD-NC to PD-MCI conversion. We employed survival analysis models on this longitudinal dataset with 17 candidate variables: 5 ThT features, 7 DLS features, neurotoxicity, and 4 demographic features (age, sex, education, and disease duration). The analysis utilized a time-varying covariate approach with a counting process data structure, allowing biomarker values to update at each follow-up visit. Four models were evaluated: Time-varying Cox for statistical inference, and LASSO-Cox, Random Survival Forest (RSF), and Gradient Boosting Survival Analysis (GBM) for prediction modeling (**Fig. 6a**).

**Figure 6.**
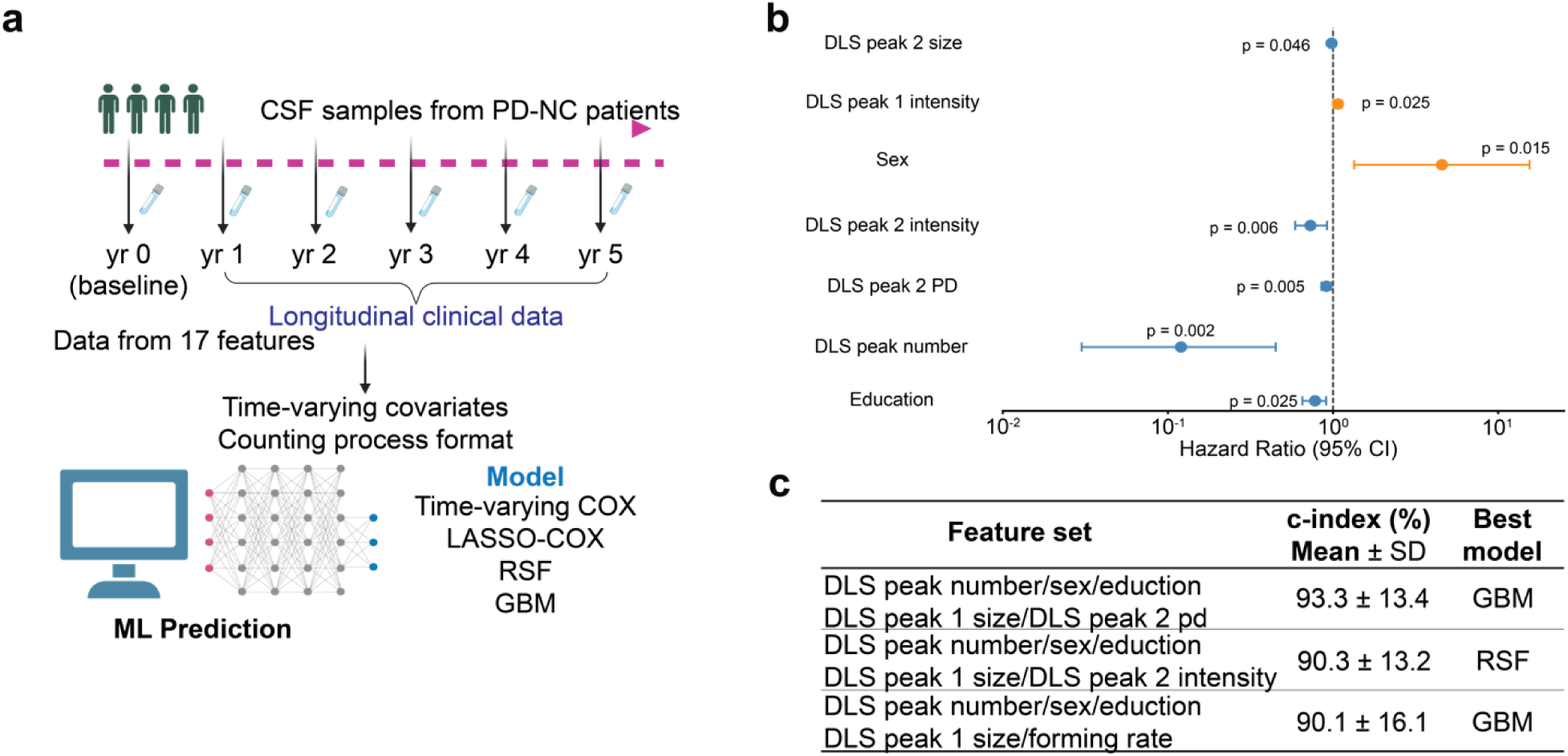
Survival analysis using ML. (**a**) Study design using longitudinal biomarker data from PD-NC patients. CSF samples were collected at baseline and follow-up visits (yr 0-5). Data from 17 features were structured in time-varying covariate counting process format and analyzed using Time-varying Cox Proportional Hazards Model, LASSO-Cox, Random Survival Forest (RSF), and Gradient Boosting Survival Analysis (GBM) models. (**b**) Forest plot of univariate time-varying Cox regression results. Hazard ratios (HR) with 95% confidence intervals are shown for significant biomarkers (p < 0.05). Blue points indicate HR < 1 (protective), orange points indicate HR > 1 (risk factor). **(C)** Prediction performance of optimal biomarker combinations. C-index values (mean ± SD) from repeated cross-validation (5-fold × 10 iterations) are shown.

**Figure 7:**
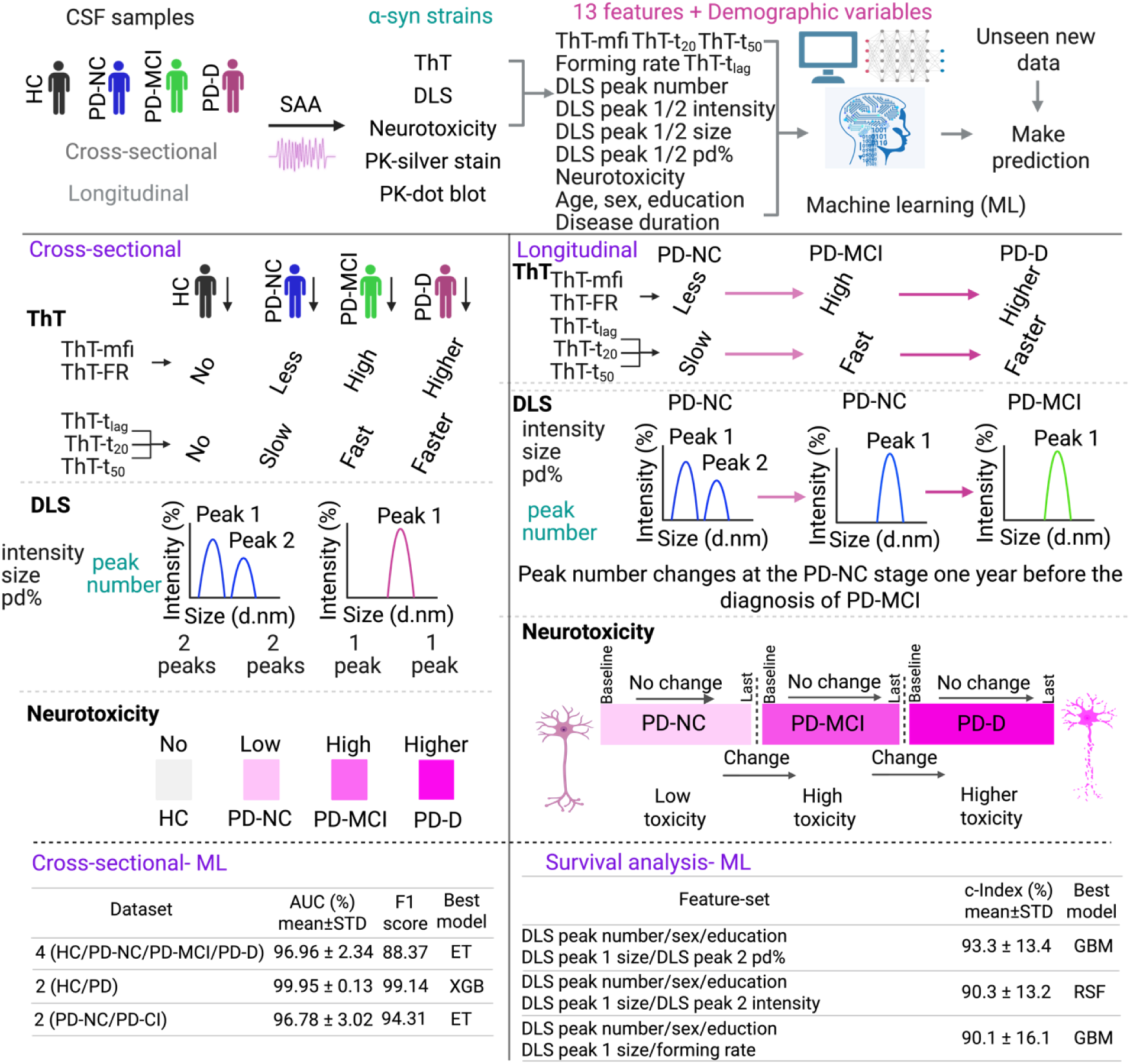
Schematic representation of α-syn strains for diagnosis and prediction of cognitive impairment in Parkinson’s disease. Both cross-sectional and longitudinal samples were used to amplify α-syn strains from CSF samples. Thioflavin T (ThT), Dynamic light scattering (DLS), neurotoxicity assays, and demographic variables yielded 17 features for disease prediction using machine learning (ML). These combined α-syn strain features provide enhanced prediction and segregation of PD subtypes and cognitive status.

Univariate time-varying Cox regression identified DLS peak number as the strongest predictor of MCI conversion (HR = 0.12, 95% CI: 0.03-0.45, *P* = 0.002), indicating that patients with a higher peak number (2 vs 1) had 88% reduced risk of cognitive decline. Sex was also significant (HR = 4.56, *P* = 0.015), with males showing ∼4.5-fold higher risk. Additional significant biomarkers included DLS peak 2 PD (HR = 0.91, *P* = 0.005), DLS peak 2 intensity (HR = 0.73, *P* = 0.006), DLS peak 1 intensity (HR = 1.07, *P* = 0.025), education (HR = 0.80, *P* = 0.025), and DLS peak 2 size (HR = 0.98, *P* = 0.046) (**Fig. 6b, Supplementary Table 23**).

Variables combinations substantially outperformed individual predictors, with optimal performance. For prediction modeling, we performed systematic forward selection by adding variables to the core variables (DLS peak number and sex), which were identified via univariate Cox regression, using repeated stratified cross-validation (5-fold × 10 iterations). Variable combinations substantially outperformed individual predictors, with optimal performance achieved using 5 variables. The best model (GBM: DLS-peak-number + sex + education + DLS-peak-1-size + DLS-peak-2-pd%) achieved 93.27% ± 13.43% C-index, while the RSF model with the same variables achieved 89.12% ± 15.59%. The 4 variable combination (GBM: DLS-peak-number + sex + education + DLS-peak-1-size) exhibited a slightly lower C-index (90.54% ± 13.49%), and the 3 variable combination (GBM: DLS-peak-number + sex + education) showed a C-index of 88.38% ± 16.46%. Six variable combinations showed decreased performance across all models, indicating overfitting and validating 5 variables as optimal complexity **(Fig. 6c, Supplementary Tables 16-21**).

DLS peak number and sex formed the backbone of top-performing models, with education, DLS peak 1 size and DLS peak 2 PD providing complementary demographic and structural information. GBM models excelled in capturing complex feature interactions, achieving the highest C-index (93.27%), while RSF also showed robust performance (89.12%). The systematic performance evolution, averaged across three machine learning models, showed progressive improvement from 2-variable (75.78%), to 3-variable (83.14%), to 4-variable (84.33%), to 5-variable (85.88%), followed by decline at 6-variable (85.41%), establishing the optimal biomarker panel that balances predictive accuracy with model parsimony. SHAP analysis of the top models revealed that DLS peak 1 size, DLS peak 2 polydispersity, and sex were the most influential features driving GBM predictions, while RSF showed more balanced contributions across all five variables (**Supplementary Table 22**).

### α-Synuclein strains become more neurotoxic with cognitive decline, but not disease duration

Using longitudinal CSF samples from Cohort I, we performed neurotoxicity assays on amplified α-syn strains collected across yearly follow-up visits. A significant increase in neurotoxicity was observed in patients transitioning to different cognitive states, specifically the PD-NC→PD-MCI, and PD-MCI→PD-D groups (**Extended Data figure 10a**). In contrast, groups without cognitive change (HC→HC, PD-NC→PD-NC, PD-MCI→PD-MCI and PD-D→PD-D), showed no significant changes in neurotoxicity during the follow-up time (**Extended Data figure 10a**). The neurotoxicity of these α-syn strains amplified from yearly collected CSF was also assessed. Notably, in the groups of HC→HC, PD-NC→PD-NC, PD-MCI→PD-MCI and PD-D→PD-D, the results demonstrated consistent neurotoxicity levels annually (**Extended Data figure 10b,c,e**). During the progression from the PD-NC to PD-MCI, and PD-MCI to PD-D, the enhanced neurotoxicity occurred at the time of the diagnosis of the cognitive change, but the neurotoxicity did not predate the cognitive decline (**Extended Data figure 10d,f,g**). Taken together, these results indicate that the neurotoxicity of amplified α-syn strains increases when PD patients undergo cognitive decline, but remains the same when the cognitive status is stable.

### Increased resistance to PK digestion when cognition progresses from PD-NC to PD-MCI and PD-D

PK digestion was performed on the amplified ɑ-syn strains over time in the same individual (**Extended Data figure 11a**). Within the groups without cognitive change, HC→HC, PD-NC→PD-NC, PD-MCI→PD-MCI and PD-D→PD-D, there was no significant difference in the remaining α-syn signal between the first and the last visits (**Extended Data figure 11a,b**). In the PD-NC→PD-MCI group, the amplified α-syn strains at the PD-MCI stage were significantly more resistant to PK digestion than PD-NC strains in the same individuals (**Extended Data figure 11a,b**). In the PD-MCI→PD-D group, PD-D strains were significantly more resistant than PD-MCI strains in the same individuals (**Extended Data figure 11a,b**). The PK digestion pattern remained stable in patients without cognitive decline (**Extended Data figure 11c,d,g**). Resistance to PK digestion was observed in patients with cognitive change: PD-NC→PD-MCI, PD-MCI→PD-D (**Extended Data figure 11e,f**). Resistance to PK digestion did not change with disease duration.

## Discussion

Our work addresses three complementary gaps in the current α-syn SAA literature on PD cognition through three integrated innovations. First, most prior studies have relied on single, baseline biomarker measurements and treated α-syn SAA readouts as time-invariant, limiting their ability to capture dynamic changes that precede cognitive decline; we overcome this by implementing a time-varying covariate survival analysis using a counting process framework, enabling longitudinal modeling of biomarker trajectories and demonstrating that specific α-syn strain features predict transition from PD with normal cognition (PD-NC) to mild cognitive impairment (PD-MCI). Second, earlier work has largely provided descriptive comparisons of a limited subset of SAA kinetic parameters, without systematically integrating structural, neurotoxic, and protease-resistant properties of α-syn strains; we address this by establishing a multi-parametric strain characterization platform that combines ThT kinetics, DLS architecture, neuronal toxicity, and proteinase K resistance to define stage-specific and within-individual strain signatures across PD-NC, PD-MCI, and PD-D. Third, existing studies have focused predominantly on diagnostic discrimination using individual biomarkers or low-dimensional models, with little emphasis on prognostic, multi-feature modeling in clinically characterized PD cohorts; we advance the field by integrating multi-dimensional strain features with demographic variables in machine-learning frameworks, revealing that synergistic combinations of a small panel of features provide high-accuracy cross-sectional classification and robust prediction of imminent cognitive decline. Longitudinal analyses further demonstrated that α-syn strain characteristics can predict cognitive deterioration in PD-NC patients up to one year before PD-MCI diagnosis. Among all features, DLS peak number showed the earliest and most reproducible change, shifting from a two-peak to a one-peak profile approximately one year before clinical PD-MCI onset, accompanied by concurrent changes in ThT kinetics, neurotoxicity, and protease resistance. Univariate time-varying Cox regression identified DLS peak number as the strongest independent predictor of cognitive transition (HR = 0.12, 95% CI: 0.03-0.45, *P* = 0.002), indicating that patients maintaining a two-peak profile had 88% reduced risk of MCI conversion. The optimal five-variable panel, consisting of DLS peak number, sex, education, DLS peak 1 size, and DLS peak 2 polydispersity, achieved a ∼93% concordance index in machine learning survival analysis using gradient boosting survival analysis (GBM). Averaged across all three machine learning models (LASSO-Cox, RSF, GBM), the five-variable panel (C-index = 85.9%) outperformed the core two-variable model (C-index = 75.8%) by 10.1% points. Notably, our longitudinal data establish a clear temporal hierarchy among biomarker modalities: DLS peak number is the only feature that changes approximately one year before clinical PD-MCI diagnosis, serving as a genuinely predictive biomarker, whereas ThT kinetics (mfi, t_lag_, t_50_), neurotoxicity, and proteinase K resistance change at the time of cognitive transition, functioning as concurrent indicators of cognitive status rather than advance predictors. This multi-parametric approach, which integrates multiple DLS features and demographic predictors into a predictive framework, represents a shift from traditional binary SAA outputs toward quantitative and prognostically informative assessments^15–17^.

Although previous studies have demonstrated that α-syn SAA can differentiate cognitive stages in PD and detect stage-dependent kinetic and structural alterations in CSF and plasma^6–10^, these investigations have largely remained descriptive and have relied on binary classifications without fully exploiting the mechanistic and prognostic potential of multi-parametric strain characterization. The present study extends this literature by integrating quantitative biophysical, biochemical, and neurotoxic properties of α-syn strains into a unified analytical framework capable of forecasting individual cognitive trajectories. Earlier reports showed that kinetic parameters such as lag phase and fluorescence amplitude, along with structural strain features, differ between PD-NC and PD-D and may precede cognitive deterioration ^6–10^. By embedding these established strain-level differences within a multi-dimensional feature space, our approach resolves subtle conformational transitions that conventional SAA cannot detect. Within this framework, DLS peak number emerged as a particularly sensitive early biomarker of conformational evolution.

The integration of machine learning with α-syn SAA represents a methodological advancement that moves the field from categorical stage discrimination toward quantitative, prognostically informative modeling of disease progression. Prior SAA studies achieved high diagnostic accuracy for distinguishing synucleinopathies, with performance values of sensitivity 86–96% and specificity 93–100%^16,18^, yet these analyses predominantly generated binary outputs with limited predictive value. Recent work demonstrated that kinetic parameters such as t₅₀ or AUC can predict phenoconversion in prodromal PD^19^, and longitudinal studies showed that aggregation and structural strain signatures can emerge up to one year before clinical PD-D onset ^6–10^. However, no previous investigation has systematically integrated multi-parametric α-syn strain features with predictive modeling in established PD cohorts. By combining ThT kinetic profiles, DLS-derived architectural characteristics, neurotoxicity measures, and protease resistance patterns within a machine learning framework, our study provides a mechanistically grounded and data-driven model of cognitive progression. This analytical shift supports the development of next-generation precision biomarkers that enable clinically actionable prognostic applications.

We achieved exceptional discriminatory performance across multiple classification tasks. In cross-sectional analysis, our 13-feature Extra Trees model distinguished four cognitive classes (HC vs. PD-NC vs. PD-MCI vs. PD-D) with 97.0% AUC, 89.2% accuracy, and 88.4% F1 score. The binary HC vs. PD classification reached 99.9% AUC (XGBoost, 13-feature set), while PD-NC vs. cognitive impairment classification achieved 96.8% AUC (Extra Trees, 17-feature set). These cross-sectional performances substantially exceed conventional clinical predictors for PD cognitive stratification, which typically achieve AUC values of 0.75–0.84^20,21^. Leave-One-Cohort-Out validation demonstrated robust generalizability across the Cohort I and Cohort II cohorts, with HC vs PD-D maintaining > 99% AUC in both directions and 4-class achieving 93.2-99.5% AUC (**Supplementary Table 15**). Importantly, comparison of biomarker-only models (13 features) with models incorporating additional demographic variables (16–17 features) revealed negligible performance differences: AUC values were identical for HC vs. PD (99.94% vs. 99.94%), and differed by only 0.80 percentage points for PD-NC vs. PD-CI (95.92% vs. 96.72%) and 0.17 percentage points for 4-class classification (96.62% vs. 96.79%) (Supplementary Tables S4, S8, S12). Consistent with these findings, feature importance analysis ranked all demographic variables in the bottom tier across all three tasks—sex, age, and education each contributed less than 1% of total importance in HC vs. PD classification, and less than 3.1% individually in the remaining tasks (Supplementary Tables S3, S7, S11). These results indicate that classification performance is predominantly driven by α-syn strain biomarkers, with demographic variables serving an auxiliary role, thereby confirming that the observed discriminatory capacity reflects genuine biophysical differences in α-syn strains across cognitive stages rather than demographic confounding.

The contrasting role of education across our analytical frameworks merits discussion. In cross-sectional classification, where the objective is to distinguish existing cognitive groups based on concurrent biomarker profiles, the pronounced biophysical differences among α-syn strains across HC, PD-NC, PD-MCI, and PD-D render demographic variables largely redundant. However, in longitudinal survival analysis, the objective shifts to predicting *when* cognitively normal PD patients will transition to MCI—a task that requires modeling individual variation in the timing of cognitive decline. In this context, education, as a proxy for cognitive reserve, becomes a critical moderating variable: patients with greater cognitive reserve can tolerate a higher burden of neuropathological change before manifesting clinical cognitive impairment. This is reflected in the univariate Cox regression, where each additional year of education was associated with a 20% reduction in MCI conversion risk (HR = 0.802, *P* = 0.025), and in the forward selection results, where the addition of education to the core model (DLS peak number + sex) produced the largest single-step improvement in averaged C-index (+7.35 percentage points). Thus, education does not confound the biomarker signal but rather calibrates its prognostic interpretation—two patients exhibiting identical DLS peak number transitions may convert at different rates depending on their cognitive reserve capacity.

Critically, in our longitudinal survival analysis using time-varying covariates, the optimal five-variable GBM achieved 93.27% ± 13.43% concordance index when predicting progression from PD-NC to PD-MCI. Multivariate Cox regression confirmed DLS peak number remained significant after adjusting for sex (HR = 0.16, *P* = 0.011). Systematic evaluation of variable combinations revealed that six-variable models showed decreased performance across all algorithms (LASSO-Cox, RSF, GBM), validating five variables as the optimal complexity and demonstrating appropriate overfitting control. This level of predictive accuracy, combined with identification of specific structural biomarkers in clinically stable PD-NC patients, represents a meaningful advance for clinical implementation.

The paradigm shift from diagnostic to prognostic application carries immediate clinical implications. Historically, α-syn biomarker studies have functioned as diagnostic adjuncts—confirmatory tests applied after cognitive impairment symptoms have prompted clinical evaluation. Our results invert this model: α-syn strain analysis now enables prospective identification of asymptomatic PD-NC patients at high risk for imminent cognitive decline, during the therapeutic window before clinical diagnosis. Imagine a clinical workflow: a 62-year-old male PD-NC patient with normal Montreal Cognitive Assessment (MoCA) score undergoes routine CSF biomarker collection and strain analysis, revealing a DLS peak number transition from 2 to 1, increased DLS peak 1 size, and decreased DLS peak 2 polydispersity. Our predictive model, incorporating these five features (including sex and education level), classifies this patient as high-risk for MCI conversion. The treating neurologist can now implement interventions—cognitive rehabilitation protocols, blood pressure optimization, sleep and cardiovascular risk factor management, and potentially enrollment in disease-modifying trials—before irreversible cognitive decline occurs. This one-year advance notice, guided by molecular biomarkers and demographic factors rather than symptom emergence, exemplifies precision medicine for neurodegeneration: early risk stratification enabling preemptive intervention. Importantly, unlike broader population screening studies, our model identifies PD-NC patients specifically—the clinically relevant population for cognitive monitoring—rather than requiring screening of asymptomatic at-risk individuals.

Several limitations warrant acknowledgment. Our longitudinal survival analysis was conducted in 21 patients (13 converters to MCI, 8 stable PD-NC patients), which is a modest cohort. However, we applied repeated stratified k-fold cross-validation (5 folds × 10 iterations) with patient-level grouping to prevent data leakage from repeated measurements. The consistent decrease in performance when adding a sixth variable across all three machine learning algorithms (LASSO-Cox, RSF, GBM) demonstrates appropriate model complexity selection despite the limited sample size. Additionally, we observed longitudinal consistency when analyzing cross-sectional data across two independent cohorts, supporting reproducibility. Leave-One-Cohort-Out validation further confirmed cross-cohort generalizability for all three classification tasks (**Supplementary Table 15**). Our cohort combined two sites (Cohort I and Cohort II) rather than employing strictly independent external validation; future studies should prospectively validate findings in separate discovery and validation cohorts. CSF collection via lumbar puncture limits repeated sampling and clinical accessibility; future efforts should establish SAA assays in blood or other peripheral tissues where preliminary evidence suggests diagnostic promise. PMCA requires specialized equipment, multi-day incubation (typically 3–5 days), and expertise in protocol execution, limiting immediate clinical scalability. Standardization of quantitative strain parameters (ThT kinetics, DLS profiles, neurotoxicity assessment) across laboratories remains incomplete, necessitating rigorous inter-laboratory calibration before widespread clinical adoption.

Several pathways warrant immediate investigation. Prospective external validation in independent PD cohorts with longitudinal cognitive follow-up is essential to confirm that our one-year prediction window generalizes across populations, centers, and ethnic backgrounds. Parallel development of high-throughput PMCA platforms with automated readouts and real-time multi-modal measurements (simultaneous ThT, DLS, and neurotoxicity assessment) will improve scalability. Investigation of alternative biomatrices is critical: if strain analysis can be validated in plasma, skin biopsy, olfactory mucosa, or submandibular gland, repeated monitoring becomes feasible for clinical practice. From a therapeutic perspective, strain dynamics should be evaluated as pharmacodynamic biomarkers in disease-modifying trials; shifts in DLS distributions, ThT kinetics, or neurotoxicity could provide early efficacy signals within months rather than requiring years of clinical endpoint assessment. Extending analysis beyond PD to MSA, dementia with Lewy bodies, and pure autonomic failure will clarify whether strain-cognitive associations are PD-specific or generalizable across synucleinopathies. Prodromal populations—individuals with isolated REM sleep behavior disorder or carriers of Leucine-Rich Repeat Kinase 2 or Glucocerebrosidase mutations—represent particularly relevant populations for early biomarker characterization and optimal intervention timing identification.

In conclusion, α-syn strain characterization combined with ML and multi-omics offers a powerful platform for early risk stratification, precision monitoring, and targeted intervention in PD and related α-synucleinopathies. Accessible biomatrices, longitudinal sampling, and automated assays will be crucial for clinical translation and will enable individualized disease management and identification of novel therapeutic windows.

## Methods

### Patient enrollment and biosample collection

#### Johns Hopkins University (Cohort I)

Individuals with PD and those without motoric evidence of parkinsonism (healthy controls) were enrolled at JHU as part of our participation in the NINDS Parkinson’s Disease Biomarker Program (PDBP)^22^. Inclusion criteria for participants with included meeting United Kingdom Parkinson’s Disease Society Brain Bank (UKBB) criteria for idiopathic PD, modified to allow for individuals with a family history of PD, and receiving levodopa therapy for their PD. Inclusion criteria for healthy controls included Montreal Cognitive Assessment score indicating no cognitive impairment (MoCA > 25) and the absence of any first-degree relative with parkinsonism. All participants had to agree to and be eligible for an annual lumbar puncture. All individuals underwent the PDBP standard set of motor, psychiatric, and cognitive assessments every 6 months for the first 5 years of the investigation as well as a one-time 12-month follow-up visit at the end of the investigation, and an annual lumbar puncture.

#### Pacific Udall Center (Cohort II)

Participants were drawn from two sites of the Pacific Udall Center of Excellence in Parkinson’s Disease Research (PUC): the Veterans Affairs Puget Sound Health Care System/University of Washington and Veterans Affairs Portland Health Care System/Oregon Health Sciences University. All participants met the UKBB clinical diagnostic criteria for PD. Individuals with a history of other neurologic disorders that would significantly impact cognition, e.g., large-vessel stroke or severe traumatic brain injury, were excluded. At all visits, each participant underwent a neurological examination, the Movement Disorder Society-Unified Parkinson’s Disease Rating Scale Part III Motor Examination (MDS-UPDRS III), and a detailed neuropsychological battery. The Clinical Dementia Rating was administered to assess the impact of any cognitive deficits on the participants’ ability to perform daily activities.

### Consensus Diagnostic Process

To ensure standardized and harmonized cognitive diagnoses across all study sites, joint diagnostic consensus conferences were conducted between the two PUC sites and with Johns Hopkins University. These conferences were designed to apply a uniform diagnostic framework and minimize inter-site variability. Each conference included at least two movement disorder specialists and a neuropsychologist who jointly reviewed all available clinical, cognitive, and functional information for each participant. Cognitive status was determined as PD-NC, PD-MCI, or PD-D according to established and validated diagnostic criteria^23,24^, as previously described^25^, with consistent application of the UKBB framework across Cohorts I and II. Regular consensus meetings reinforced diagnostic reliability and ensured consistent classification across cohorts and institutions.

### Expression and purification of α-synuclein protein

Recombinant human α-syn protein was prepared according to the previous method^26^. pRK172-α-syn plasmid was transduced in BL21 (DE3) cells and cultured at 37°C in lysogeny broth overnight. The E.coli pellets were resuspended with osmotic shock buffer (3.63 g Tris-base, 400 g sucrose, and 0.744 g EDTA were dissolved in 1 L DDW, pH 7.2) by drastic agitation. The mixture was then centrifuged at 10,000 g for 30 min to remove the supernatant, and DDW containing proteinase inhibitor and 80 μL saturated MgCl_2_ were added to resuspend the pallets. The supernatant was collected (10,000 g, 30 min centrifugation) and filtered through a 0.45 μm filter, followed by the dialysis with low salt buffer (20 mM Tris-base, 50 mM NaCl in DDW; pH 8.0) overnight at 4°C. α-syn protein was purified with fast protein liquid chromatography (FPLC) and saved in a −80 °C freezer. The purity was evaluated with Coomassie brilliant blue staining and immunoblot. The concentration was measured with a BCA assay.

### Amplification of pathogenic α-syn in patient CSF with SAA

The amplification of α-syn strains from patient-derived CSF samples was performed using the SAA method by referring to previous work^1,27,28^ with some modifications. The SAA equipment containing the microplate horn (#431MPX), a sound enclosure (#432MP), and a thermoelectric chiller (#4900) was purchased from Qsonica. Briefly, recombinant α-syn was centrifuged at 100,000 g for 30 min at 4°C to remove any preformed aggregates before use. Then, α-syn was diluted with SAA buffer (1% Triton X-100 in PBS), and 100 μL was transferred into PCR tubes containing a suitable amount of silicon beads (diameter 1.0 mm, purchased from BioSpec products), and 10 μL CSF samples were added as seeds in triplicate. The final concentration of α-syn was 0.3 mg/mL. After mixing, the samples were subjected to sonication (Amplitude: 5; 40 sec sonication and 29 min 20 sec incubated at 37°C). In total there were 40 amplification circles for 1-day reaction and 280 amplification circles for 7-day reaction. 5 μL samples were collected every day and amplification was monitored by measuring Thioflavin T (ThT) (Sigma-Aldrich, cat No. T3516) fluorescence using a Fluorescence Spectrophotometer (Varioskan LUX plate reader, Thermo Fisher Scientific) with fixed excitation and emission wavelength at 450 nm and 485 nm respectively. After 7 days, the mixture was transferred into the centrifugal filter (Millipore, MCW: 3000) containing 15 mL PBS, and centrifuged at 4000 g for 30 min at 4°C to remove Triton X-100. The washing with 15 mL PBS and centrifugation was repeated 8 times and the final SAA products were collected. SAA products were spun for 30 min at 20,000g, and the amount of monomeric α-syn in the supernatant was assessed by BCA assay and the pelleted assemblies were resuspended in PBS buffer. To ensure that observed differences reflect intrinsic strain properties rather than variable seeding capacity, we perform serial dilutions of all CSF-derived seeds, establishing precise limits of detection (LOD) and quantification (LOQ). Samples below the LOD are classified as negative, while those between LOD and LOQ are reported as detectable but not quantitatively reliable. Serial passaging of SAA products further reduces variability from heterogeneous seed populations and CSF matrix effects, selectively amplifying dominant strains. We quantify seeding activity using t₅₀ and integrate maximum amplification rate (Vmax) to capture both seed abundance and amplification efficiency, allowing discrimination among α-syn strains from PD-NC, PD-MCI, and PD-D. Standardized pre-analytical procedures, such as uniform dilution or ultracentrifugation, minimize matrix effects, and Z-score normalization across groups ensures robust comparisons.

### ThT fluorescence assay

The SAA sample (5 μL) was taken out and added into a 55 μL ThT solution (20 μM). Samples were subsequently plated in triplicate on 384 well black/clear bottom plates (Sigma-Aldrich, cat no. P6491), and the fluorescence was measured at 450/485 nm excitation/emission with a microplate reader (Varioskan LUX plate reader, Thermo Fisher Scientific). Following data acquisition, the kinetic curves were fitted to Equation. 1 to calculate the half-time (t_50_) value and lag time (t_lag_) for each curve.

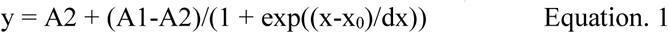

Where A1 represents initial fluorescence, A2 is the final fluorescence value, x_0_ is the half-time (t_50_) value, and dx represents the time constant.

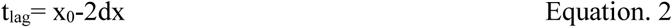

The maximum aggregation (forming) rate was determined as the first derivative of the four-parameter logistic (4-PL) fit of ThT fluorescence intensity over time, calculated at the inflection point (X0) using the equation:

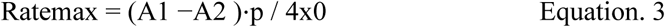

Where X0 (corresponding to the T₅₀) denotes the inflection point of the curve, and p is the slope factor describing the steepness of the aggregation phase.

### Proteinase K digestion, dot blot, and silver staining

The SAA samples (7 μg) were mixed with proteinase K (PK) and incubated at 37°C at different time points (0, 5, 15, and 30 min). For the dot blot assay, the PK-digested SAA samples were loaded onto the nitrocellulose membrane (Bio-Rad, cat no. 1620112) and blocked by 5% bovine serum albumin (BSA) (Sigma-Aldrich, cat no. A7906) in TBST for 1 hr at room temperature (RT). The membrane was then transferred into the mouse anti-α-syn mAb (1:2000 dilution, BD Biosciences, cat no. 610787) in TBST with 5% BSA overnight at 4°C. Following with TBST wash (3 times x 5 min), the membrane was incubated with anti-mouse IgG-HRP (1:5000 dilution, GE Healthcare, cat no. NA931) for 1 hr at RT. After TBST wash, the signal was developed with SuperSignal West Pico Plus chemiluminescent substrate (Thermo Fisher Scientific, cat no. 34096). For the silver staining, PK-digested SAA samples were loaded on the SDS-PAGE (15%) gels. Silver staining was performed using Pierce Silver Stain Kit (Thermo Fisher Scientific, cat no. 24612). All the images were acquired and processed with Amersham Image 600 (GE Healthcare Life Sciences).

### Dynamic light scattering (DLS)

The SAA sample (10 μg) was mixed with filtered phosphate buffer (990 μL). Measurements were performed using a Zetasizer Nano-ZS (Malvern Instruments, Malvern, UK) equipped with a He-Ne laser. Each sample was measured in 1-cm path-length polystyrene semi-micro disposable cuvettes (Fisher Emergo, Landsmeer, The Netherlands). The cell holder was maintained at 25 °C. For each sample, 10 runs were performed, with three repetitions.

### Primary cortical neuron culture, neurotoxicity of SAA products

Mouse primary cortical neurons were cultured from embryonic 15.5-day pups of CD-1 pregnant mice (Charles River). 48 well plates were coated via Poly-L-ornithine solution (0.2 mg/mL) for 1 hr at 37°C and washed 3 times with sterile dd-water. Primary neurons at 7 days in vitro (DIV) were treated with SAA samples with a final concentration of 10 μg/well. The neuropathology and neurotoxicity were assessed at 21 and 28 DIV individually. The primary cortical neurons were washed with PBS, fixed in 4% paraformaldehyde (PFA), followed by blocking in 3% goat serum containing PBST (0.1% Tween-20) for 1 hr. Anti-NeuN (1:250, MAB377, Sigma-Aldrich) were incubated overnight at 4°C, followed by Alexa-fluor 488 secondary antibodies (1:2000, Thermo Fisher Scientific) and Hoechst (1:5000, Thermo Fisher Scientific) at RT for 1 hr. The fluorescence images were obtained via a Nice microscope (Zeiss). The number of NeuN was quantified using ImageJ software (National Institute of Health, Bethesda, MD).

### Statistical analysis

Statistical analysis was performed using the statistical software Stata (version 18). The baseline demographic and clinical characteristics of the participants are presented as mean +/- standard deviation (SD) or number (%). The characteristics were compared using the Kruskal-Wallis or chi squared as appropriate. Logistic regression analyses were performed with the binary cognition variable as the outcome variable and the biomarker of interest as the independent variables, adjusting for covariates, such as age and sex. Receiver operating characteristic curves were calculated to visualize and compare the predictivity of the biomarkers. Two-sided p-values <0.05 were considered significant. For the multivariable logistic regression analyses (**Supplementary Tables 24-27**), each biomarker was entered as the independent variable with cognitive status (PD-NC vs PD-CI) as the outcome, adjusting for sex, age, education, and disease duration as covariates. These four demographic variables were selected a priori based on their established associations with cognitive decline in PD.

### Disease prediction using machine learning (ML)

#### Cross-sectional classification

We implemented machine learning classifiers for three cross-sectional classification tasks (HC vs PD, PD-NC vs PD-CI, and 4-class classification) using patient sample data. The dataset comprised 13 α-syn strain features and 4 demographic variables (age, sex, education, and disease duration). We selected six tree-based models (Catboost, Gradient Boosting Decision Trees, XGBoost, Random Forest, Decision Trees, and Extra Trees) and one linear model (Logistic Regression with L1 regularization) with default hyperparameters. For the HC vs PD-D task, disease duration was excluded from the feature set as all HC subjects have a disease duration of zero by definition, rendering this variable trivially discriminating rather than biologically informative; thus, 16 features were used for Task 1 and 17 features for Tasks 2 and 3. For feature selection and model evaluation, we employed two complementary approaches: a Top-K strategy using Random Forest feature importance rankings, and predefined feature sets based on measurement type (ThT-based, DLS-based, neurotoxicity, and combinations with demographic variables). Model performance was assessed using repeated stratified k-fold cross-validation (5-fold, 10 iterations) with AUC, accuracy, precision, recall, and F1 score. For the 4-class classification, CatBoost was excluded due to compatibility issues with multi-class settings. Generalizability was further evaluated using Leave-One-Cohort-Out validation across two independent cohorts (Cohort I and Cohort II). Full numerical results for all cross-sectional classification models, including individual feature performance, feature combinations, feature importance, confusion matrices, and classification reports, are provided in **Supplementary Tables 1–15**. More details can be found in Supplementary Methods.

#### Survival Analysis for longitudinal study

We utilized longitudinal biomarker data from PD-NC patients for training and evaluating survival analysis models to predict cognitive decline progression (PD-NC to PD-MCI). The dataset comprised 17 candidate variables (13 biomarkers, age, sex, education, and disease duration) across 44 observation intervals from 21 patients (13 converters, 8 stable). The analysis employed a time-varying covariate approach using counting process data structure (start, stop, event), which allows biomarker values to update at each follow-up visit rather than using only baseline measurements. The analysis employed a time-varying covariate approach using counting process data structure (start, stop, event), which allows biomarker values to update at each follow-up visit rather than using only baseline measurements. Three machine learning survival models were evaluated for prediction: LASSO-Cox^29^, Random Survival Forest (RSF)^30^, and Gradient Boosting Survival analysis (GBM), alongside a Time-varying Cox Proportional Hazards model for statistical inference. Core predictor variables (DLS peak number and sex) were identified through univariate Cox regression based on hazard ratio magnitude, and optimal variable combinations were determined through systematic forward selection (2 to 6 variables), with the optimal set confirmed where average C-index was maximized and subsequently declined upon adding further variables. Model performance was evaluated using repeated stratified group k-fold cross-validation (5-fold, 10 iterations) with patient-level grouping to prevent data leakage. The Concordance Index (C-index)^31^ served as the primary metric, and SHAP analysis was performed on the top three model configurations to quantify feature importance. Full numerical results for all survival analysis models are provided in **Supplementary Tables 16–23**. More details can be found in Supplementary Methods..

## Acknowledgements

The authors acknowledge Jiangxia Wang for help with the statistical analysis discussion. The Multiphoton Imaging Core of Johns Hopkins University was used (NS050274) in some of the imaging studies. T.M.D. is the Leonard and Madlyn Abramson Professor in Neurodegenerative Diseases. For the purpose of open access, the author has applied a CC-BY public copyright license to the Author Accepted Manuscript (AAM) version arising from this submission.

## Author contributions

KG performed all the basic science experiments and data analysis. KK, CHN and SZH performed data analysis using ML and conceptualization. NW, EQX, XDZ, HYL, JD, JY, AW, YJC, RK, HQL, LLN, Rong-C, and SZ contributed to biophysical, biochemical, cellular experiments and data interpretation. CB, BC, JFQ, KAC, ALH, SCH, TJM, CPZ, and LSR provided key patients’ samples and clinical information. JXW assisted with the statistical analysis. LTJ, YJL, CHN and MYY helped with the concept generation, manuscript writing, and experiment design. XM designed the project direction and research strategy. XM, JSR, TMD, VLD, and CHN wrote and revised the paper. All authors reviewed, edited, and approved the paper.

## Data and materials availability

Further information and requests for resources and reagents should be directed to and will be fulfilled by Xiaobo Mao. There are no restrictions on any data or materials presented in this paper. All data are available in the main text or the Extended Data.

---

**TOC:**
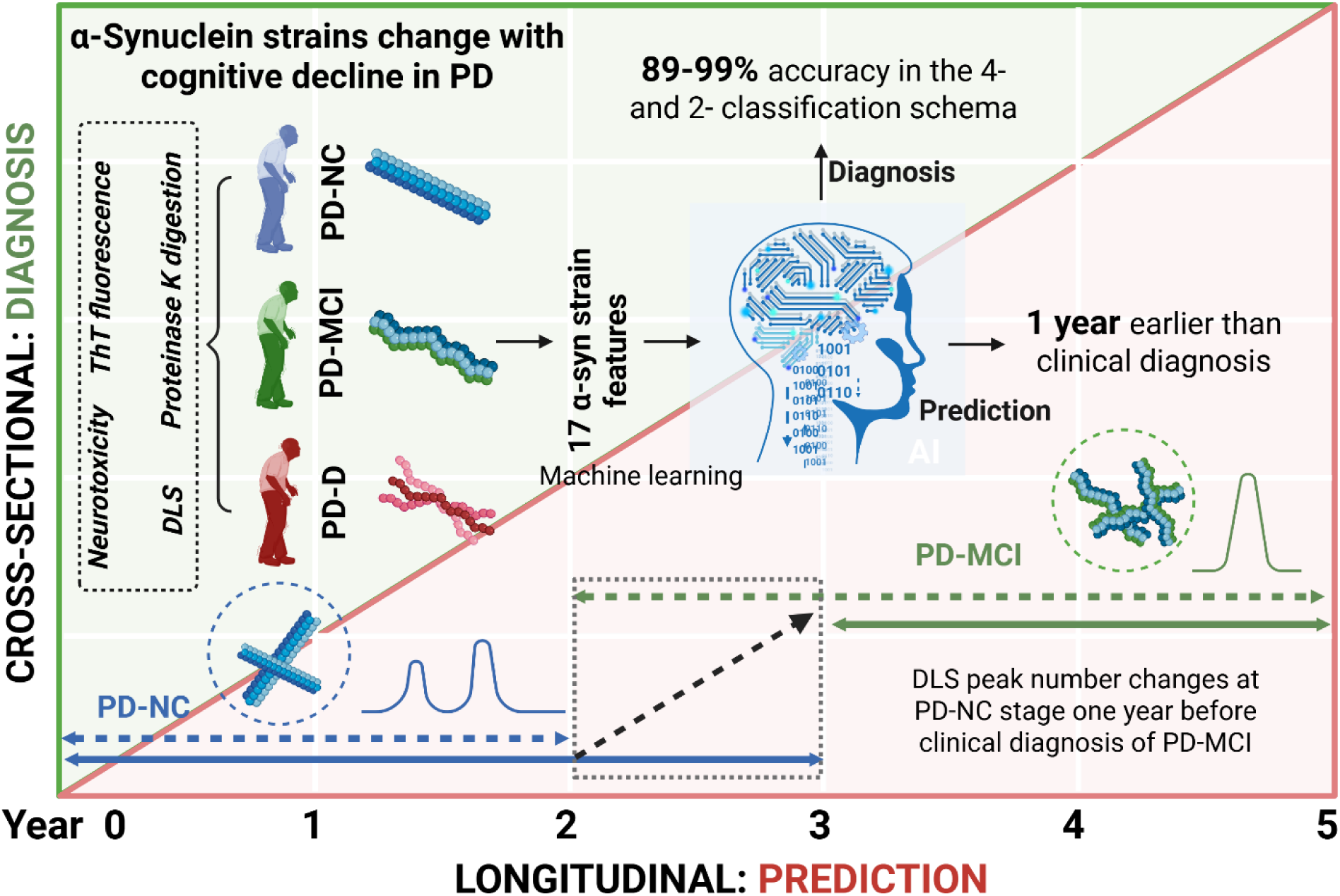
α-Synuclein strain changes allied with cognitive changes in Parkinson’s disease. Cross-sectional and longitudinal studies show that α-syn strain features (Thioflavin T (ThT), Dynamic light scattering (DLS), and neurotoxicity) change with and before cognitive decline in PD. Machine learning greatly enhances the diagnostic ability to predict PD subtypes and cognitive status.

## Supporting information

**Extended Data figure 1.**
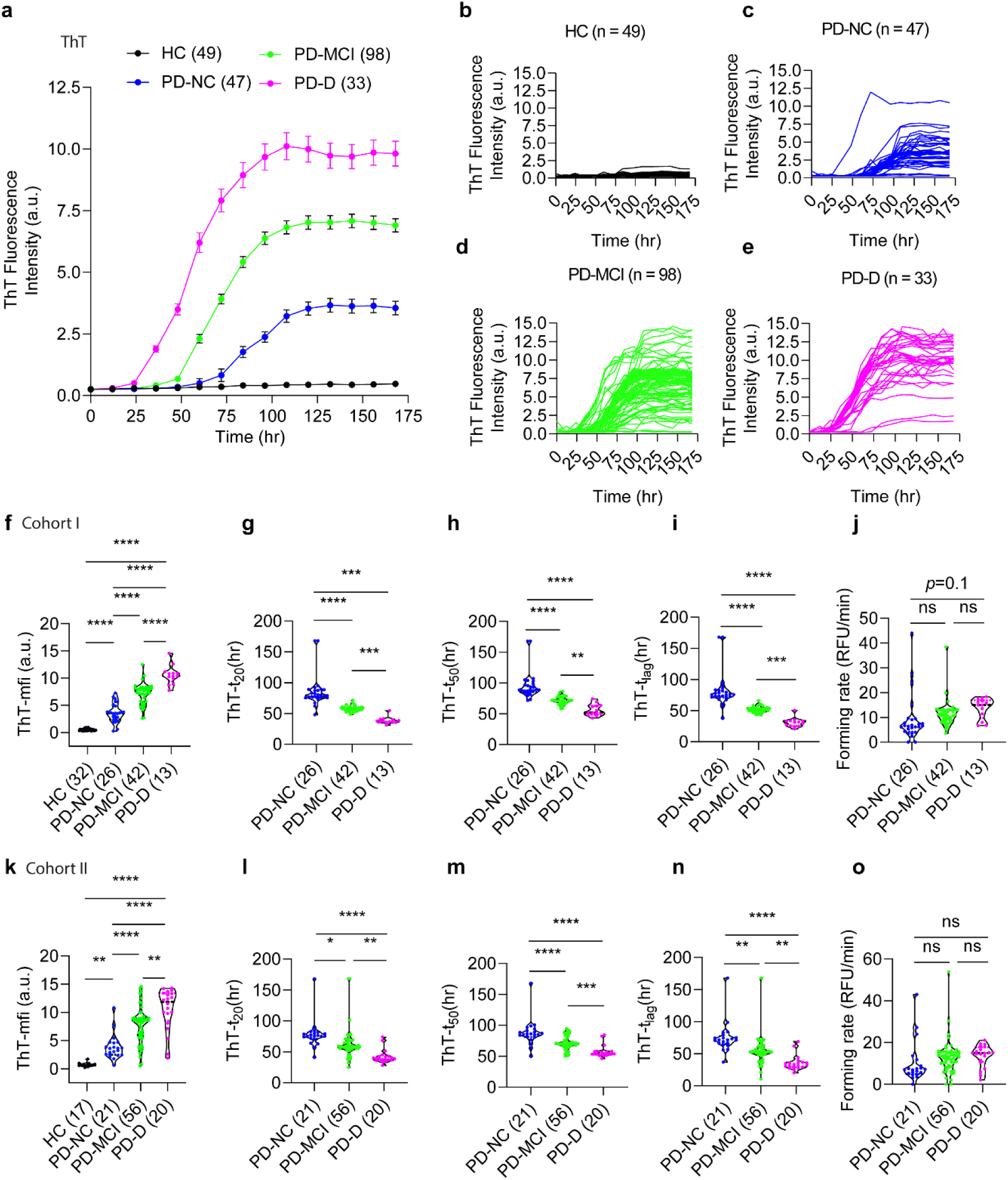
ThT assay for amplified α-syn strains from cohort I and II. (**a**) Averaged ThT fluorescence spectra for amplified α-syn strains from HC, PD-NC, PD-MCI, and PD-D. (**b, c, d,** and **e**) ThT fluorescence spectra for individual samples from HC, PD-NC, PD-MCI, and PD-D respectively. **(f-j**) ThT-mfi (**f**), ThT-t_20_ (**g**), ThT-t_50_ (**h**), ThT-t_lag_ (**i**), and forming rate (**j**) for samples from cohort I. **(k-o**) ThT-mfi (**k**), ThT-t_20_ (**l**), ThT-t_50_ (**m**), ThT-t_lag_ (**n**), and forming rate (**o**) for samples from cohort II. Data are presented as the mean ± SEM. The statistical significance was evaluated via one-way ANOVA with Tukey’s multiple comparisons test. No significant difference (ns) *P* > 0.05, \**P* < 0.05, \*\**P* < 0.01, \*\*\*\**P* < 0.0001.

**Extended Data figure 2.**
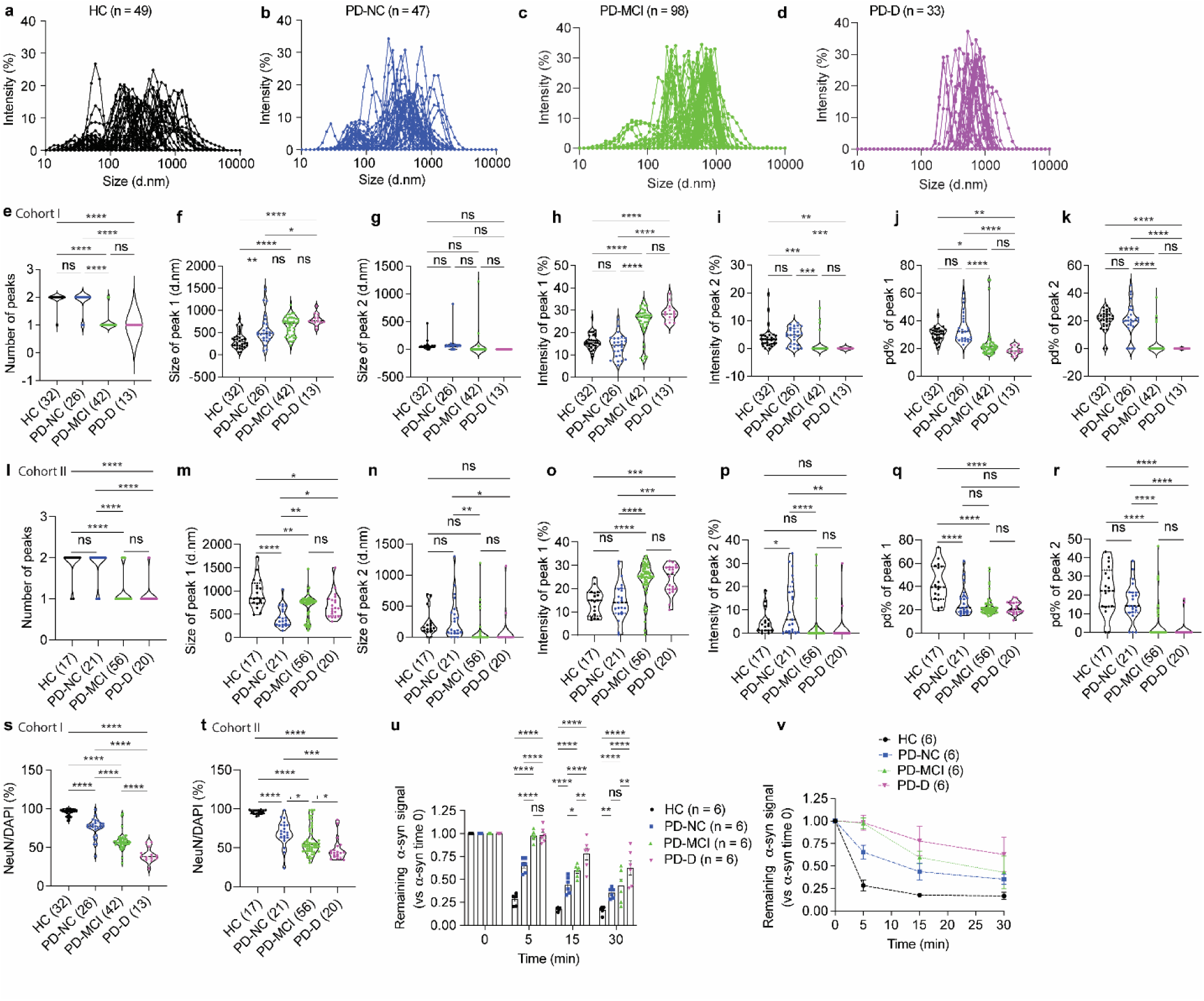
DLS and neurotoxicity of amplified α-syn strains from cohort I and II. (**a, b, c,** and **d**) DLS spectra for individual samples from HC, PD-NC, PD-MCI, and PD-D respectively. Two peaks were observed in the HC and PD-NC group and one peak in PD-MCI and PD-D. Each peak represents an individual biological sample. (**e,f,g,h,i,j,k**) DLS peak number, peak size, peak intensity and pd% of peak 1 and peak 2 for α-syn strains from HC, PD-NC, PD-MCI, and PD-D from cohort I. (**l,m,n,o,p,q,r**) DLS peak number, peak size, peak intensity and pd% of peak 1 and peak 2 for α-syn strains from HC, PD-NC, PD-MCI, and PD-D from cohort II. **(s**) Neuron viability for samples from HC, PD-NC, PD-MCI, and PD-D from cohort I. **(t**) Neuron viability for samples from HC, PD-NC, PD-MCI, and PD-D from cohort II. **(u,v**) α-syn signal remaining after PK digestion as assessed by dot-blot for α-syn immunoreactivity. Amplified α-syn strains were treated with PK and assessed at different time points (0, 5, 15, and 30 min) after PK administration. Data are presented as the mean ± SEM. The statistical significance was evaluated via one-way ANOVA with Tukey’s multiple comparisons test. No significant difference (ns) *P* > 0.05, \**P* < 0.05, \*\**P* < 0.01, \*\*\*\**P* < 0.0001.

**Extended Data figure 3.**
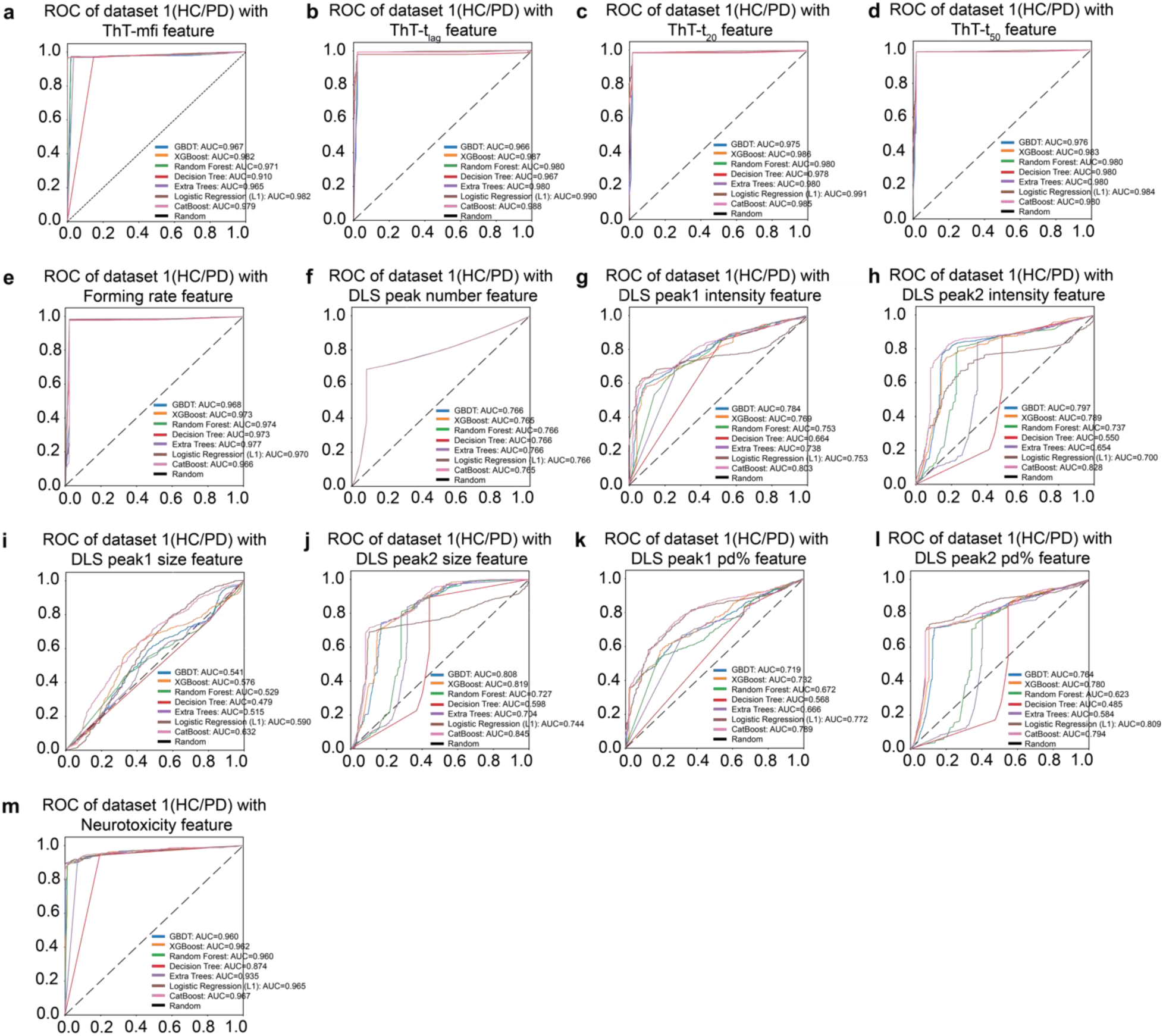
ROC of dataset 1 (HC/PD) with ThT, DLS and neurotoxicity feature using 7 models. (**a**) ROC of dataset 1 (HC/PD) with ThT-mfi feature. (**b**) ROC of dataset 1 (HC/PD) with ThT-t_lag_ feature. (**c**) ROC of dataset 1 (HC/PD) with ThT-t_20_ feature. (**d**) ROC of dataset 1 (HC/PD) with ThT-t_50_ feature. (**e**) ROC of dataset 1 (HC/PD) with forming rate feature. (**f**) ROC of dataset 1 (HC/PD) with DLS peak number feature. (**g**) ROC of dataset 1 (HC/PD) with DLS peak 1 intensity feature. (**h**) ROC of dataset 1 (HC/PD) with DLS peak 2 intensity feature. (**i**) ROC of dataset 1 (HC/PD) with DLS peak 1 size feature. (**j**) ROC of dataset 1 (HC/PD) with DLS peak 2 size feature. (**k**) ROC of dataset 1 (HC/PD) with DLS peak 1 pd% feature. (**l**) ROC of dataset 1 (HC/PD) with DLS peak 2 pd% feature. (**m**) ROC of dataset 1 (HC/PD) with neurotoxicity feature using 7 models.

**Extended Data figure 4.**
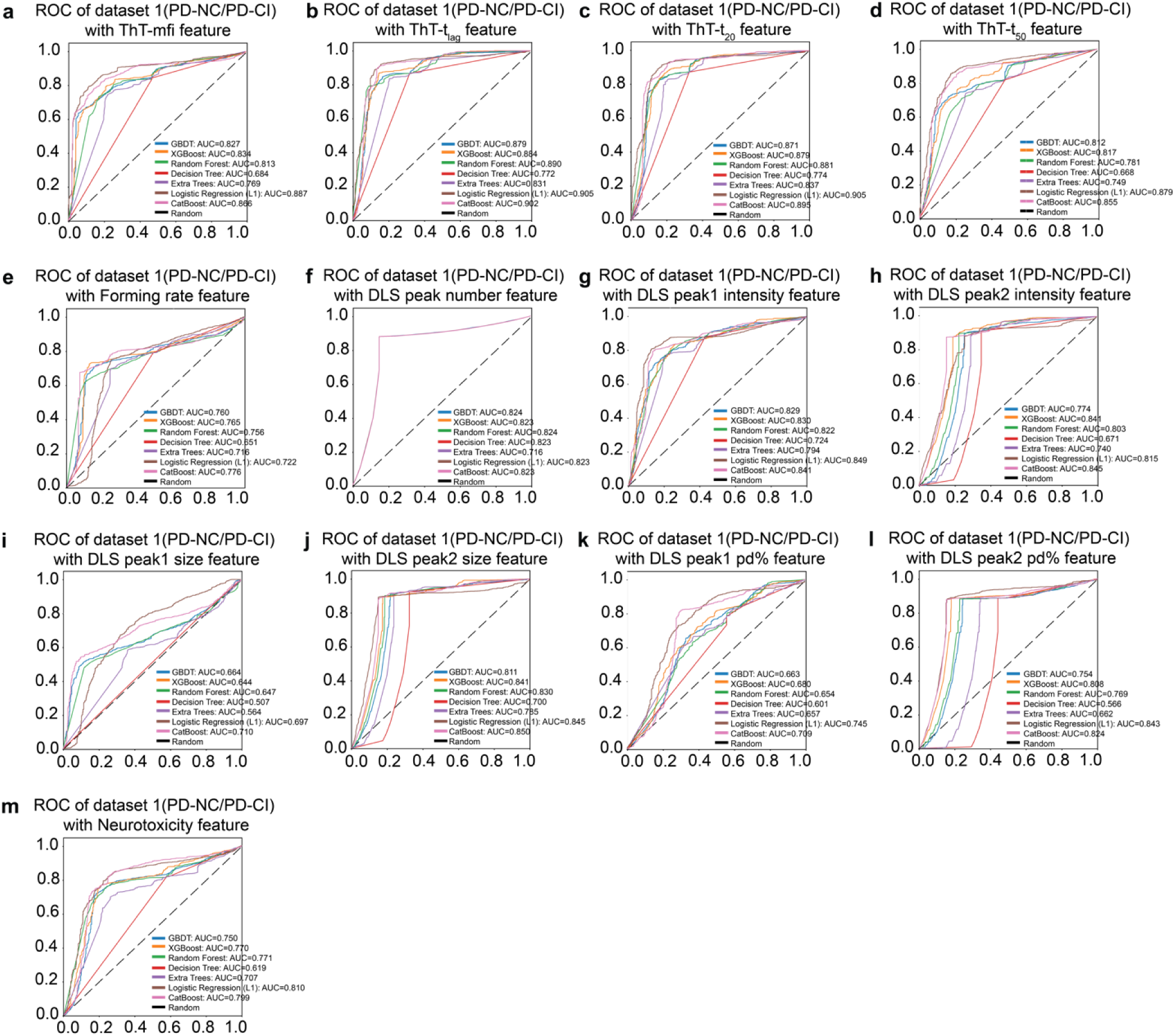
ROC of dataset 2 (PD-NC/PD-CI) with ThT, DLS, and neurotoxicity feature using 7 models. (**a**) ROC of dataset 1 (PD-NC/PD-CI) with ThT-mfi feature. (**b**) ROC of dataset 1 (PD-NC/PD-CI) with ThT-t_lag_ feature. (**c**) ROC of dataset 1 (PD-NC/PD-CI) with ThT-t_20_ feature. (**d**) ROC of dataset 1 (PD-NC/PD-CI) with ThT-t_50_ feature. (**e**) ROC of dataset 1 (PD-NC/PD-CI) with forming rate feature. (**f**) ROC of dataset 1 (PD-NC/PD-CI) with DLS peak number feature. (**g**) ROC of dataset 1 (PD-NC/PD-CI) with DLS peak 1 intensity feature. (**h**) ROC of dataset 1 (PD-NC/PD-CI) with DLS peak 2 intensity feature. (**i**) ROC of dataset 1 (PD-NC/PD-CI) with DLS peak 1 size feature. (**j**) ROC of dataset 1 (PD-NC/PD-CI) with DLS peak 2 size feature. (**k**) ROC of dataset 1 (PD-NC/PD-CI) with DLS peak 1 pd% feature. (**l**) ROC of dataset 1 (PD-NC/PD-CI) with DLS peak 2 pd% feature. (**m**) ROC of dataset 1 (PD-NC/PD-CI) with neurotoxicity feature using 7 models.

**Extended Data figure 5.**
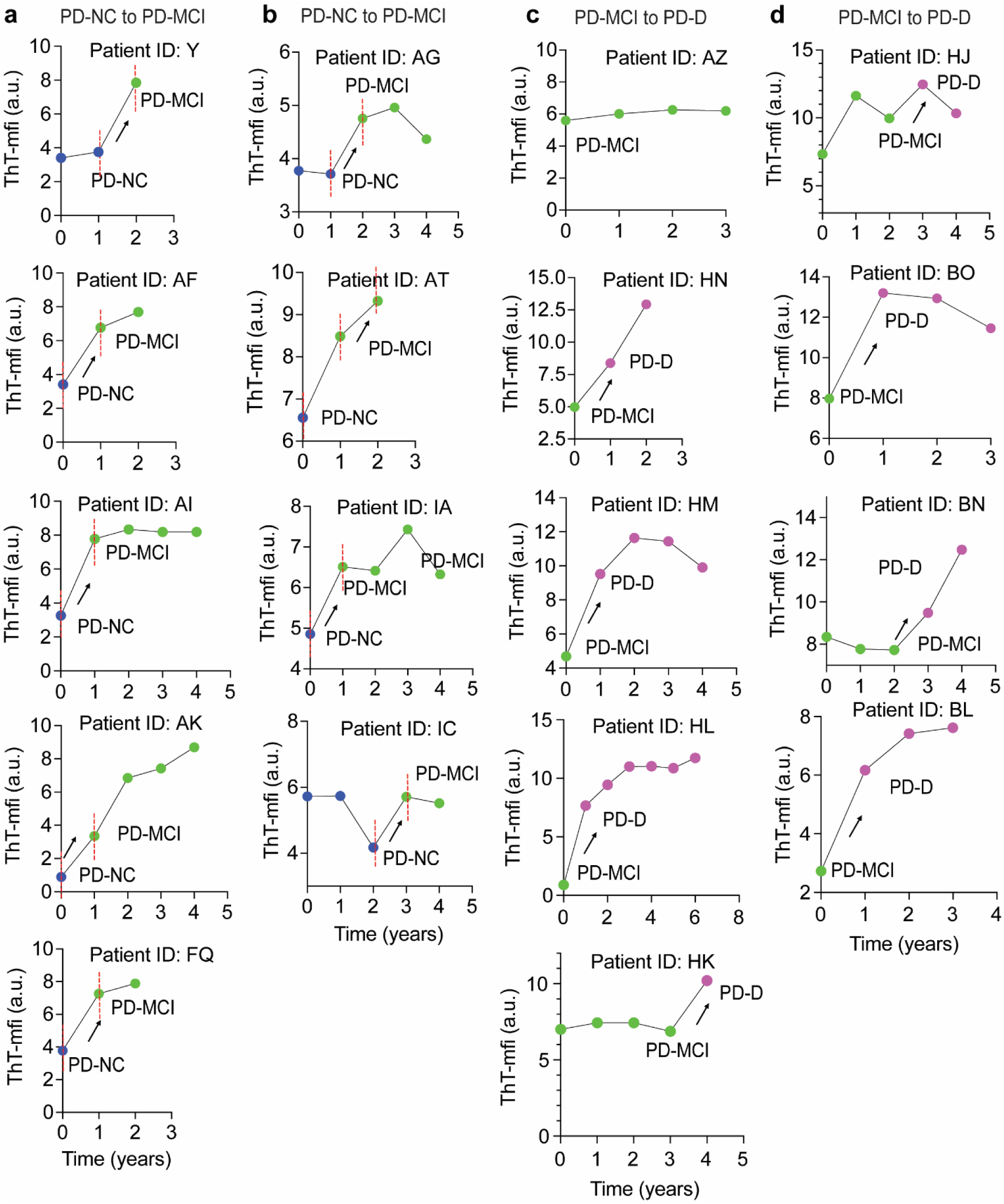
Longitudinal analysis of α-syn strains using ThT-mfi. ThT-mfi of amplified α-syn strains from HC, PD-NC, PD-MCI, and PD-D groups. **(a,b,c,d)** Yearly mapping of ThT-mfi of amplified α-syn strains from individuals with changed cognitive status.

**Extended Data figure 6.**
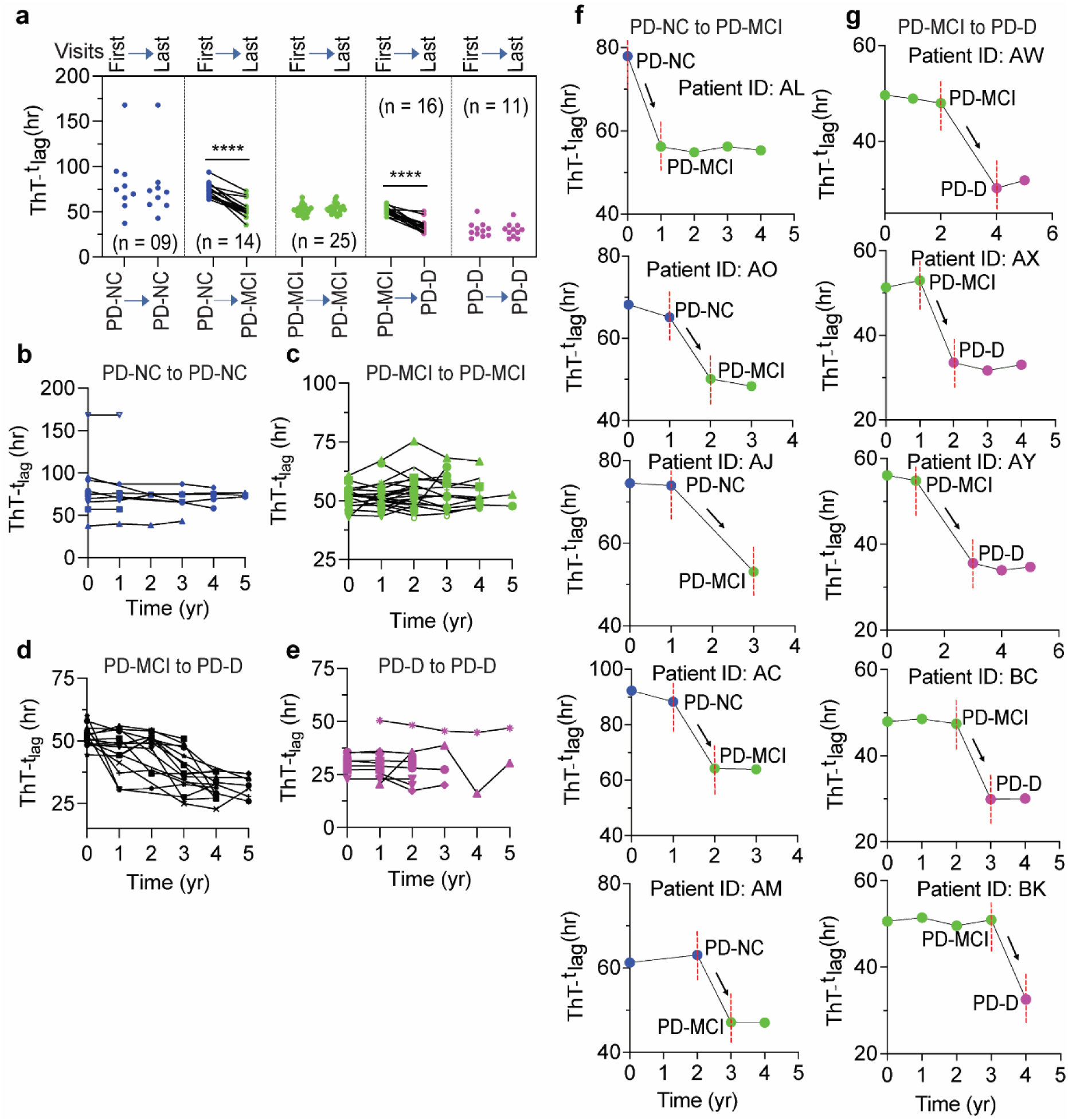
Longitudinal analysis (yearly) for α-syn strains using ThT-t_lag_. (**a**) ThT-t_lag_ of amplified α-syn strains from HC, PD-NC, PD-MCI, and PD-D groups between the first- and last-visit in Cohort I. **(b-e)** Yearly mapping of ThT-t_lag_ of amplified α-syn strains from individuals with stable cognitive status. **(f,g)** Yearly mapping of ThT-t_lag_ of amplified α-syn strains from individuals with changed cognitive status. The statistical significance was evaluated via one-way ANOVA with Tukey’s multiple comparisons test. No significant difference (ns) *P* > 0.05, \**P* < 0.05.

**Extended Data figure 7.**
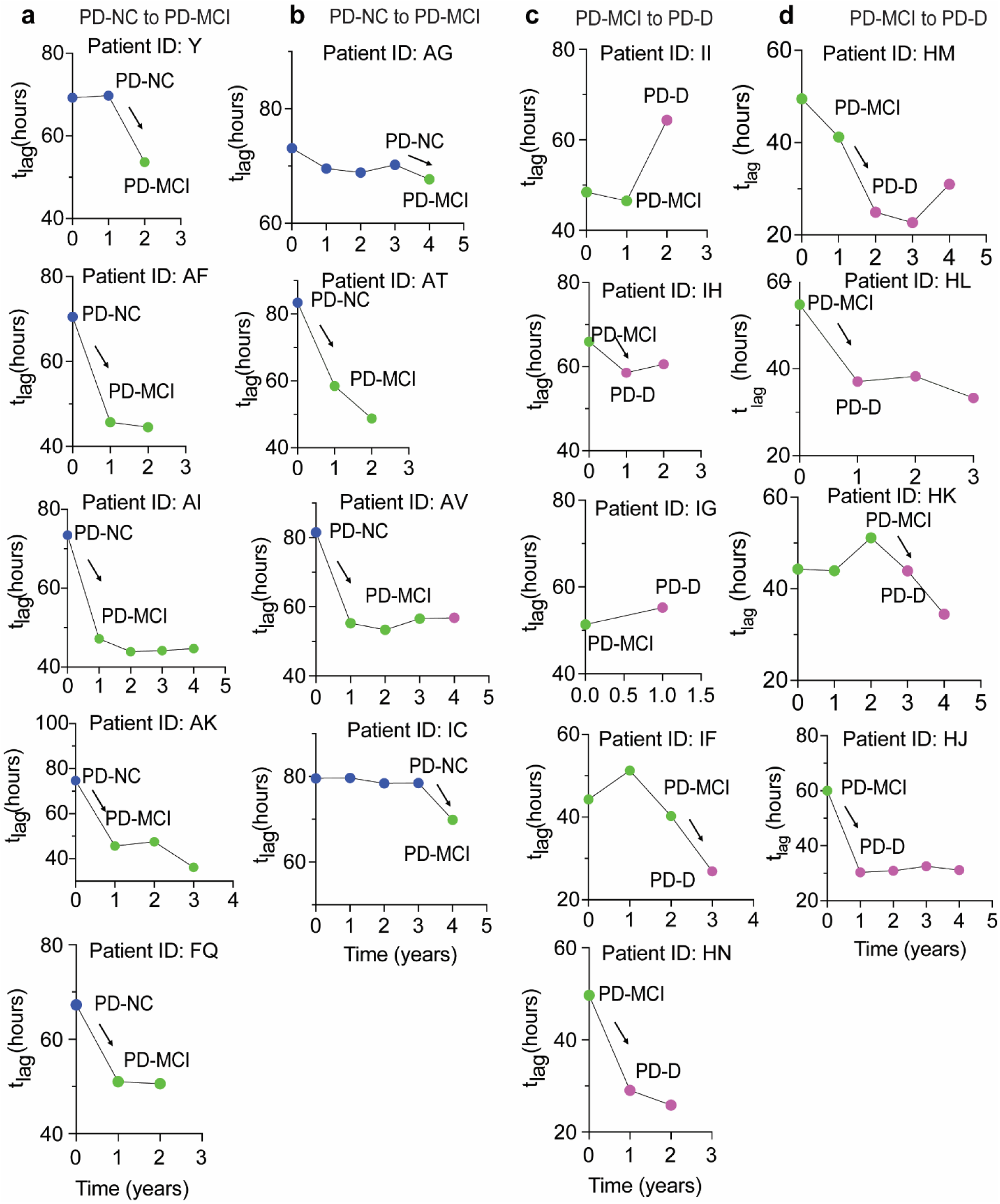
Longitudinal analysis (yearly) for α-syn strains using ThT-t_lag_. ThT-t_lag_ of amplified α-syn strains from HC, PD-NC, PD-MCI, and PD-D groups. **(a,b,c,d)** Yearly mapping of ThT-t_lag_ of amplified α-syn strains from individuals with changed cognitive status.

**Extended Data figure 8.**
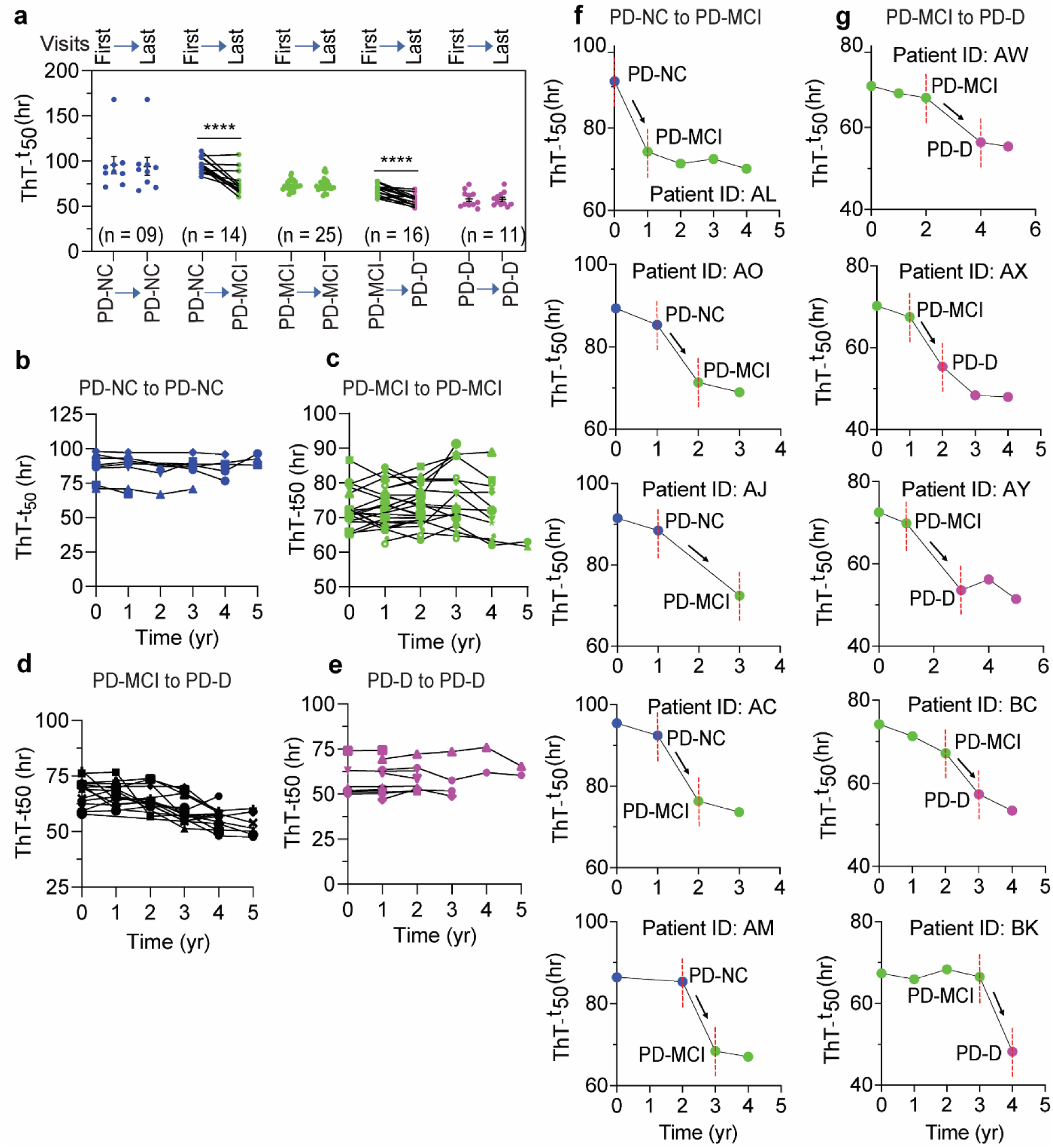
Longitudinal analysis (yearly) for α-syn strains using ThT-t_50_. (**a**) ThT-t_50_ of amplified α-syn strains from longitudinal HC, PD-NC, PD-MCI, and PD-D groups between the first- and last-visit in Cohort I. **(b,e)** Yearly mapping of ThT-t_50_ of amplified α-syn strains from individuals with stable cognitive status. **(c,d)** Yearly mapping of ThT-t_50_ of amplified α-syn strains from individuals with changed cognitive status. The statistical significance was evaluated via one-way ANOVA with Tukey’s multiple comparisons test. No significant difference (ns) *P* > 0.05, \*\*\*\**P* < 0.0001.

**Extended Data figure 9.**
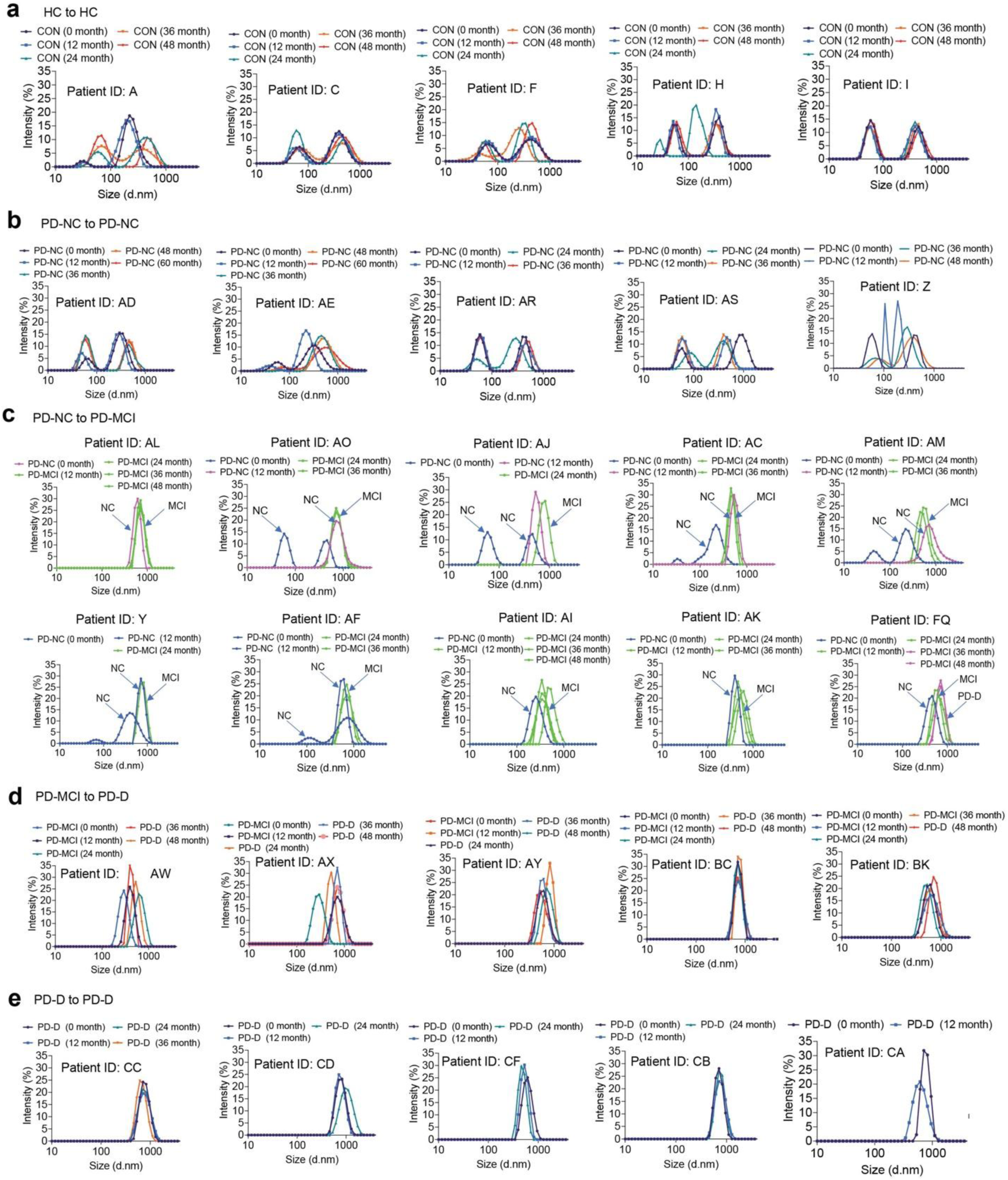
Longitudinal analysis (yearly) for α-syn strains from HC, PD-NC, PD-MCI, and PD-D with DLS. **(a)** HC to HC; **(b)** PD-NC to PD-NC; **(c)** PD-NC to PD-MCI; **(d)** PD-MCI to PD-D; **(e)** PD-D to PD-D. Each peak represents an individual biological sample.

**Extended Data figure 10.**
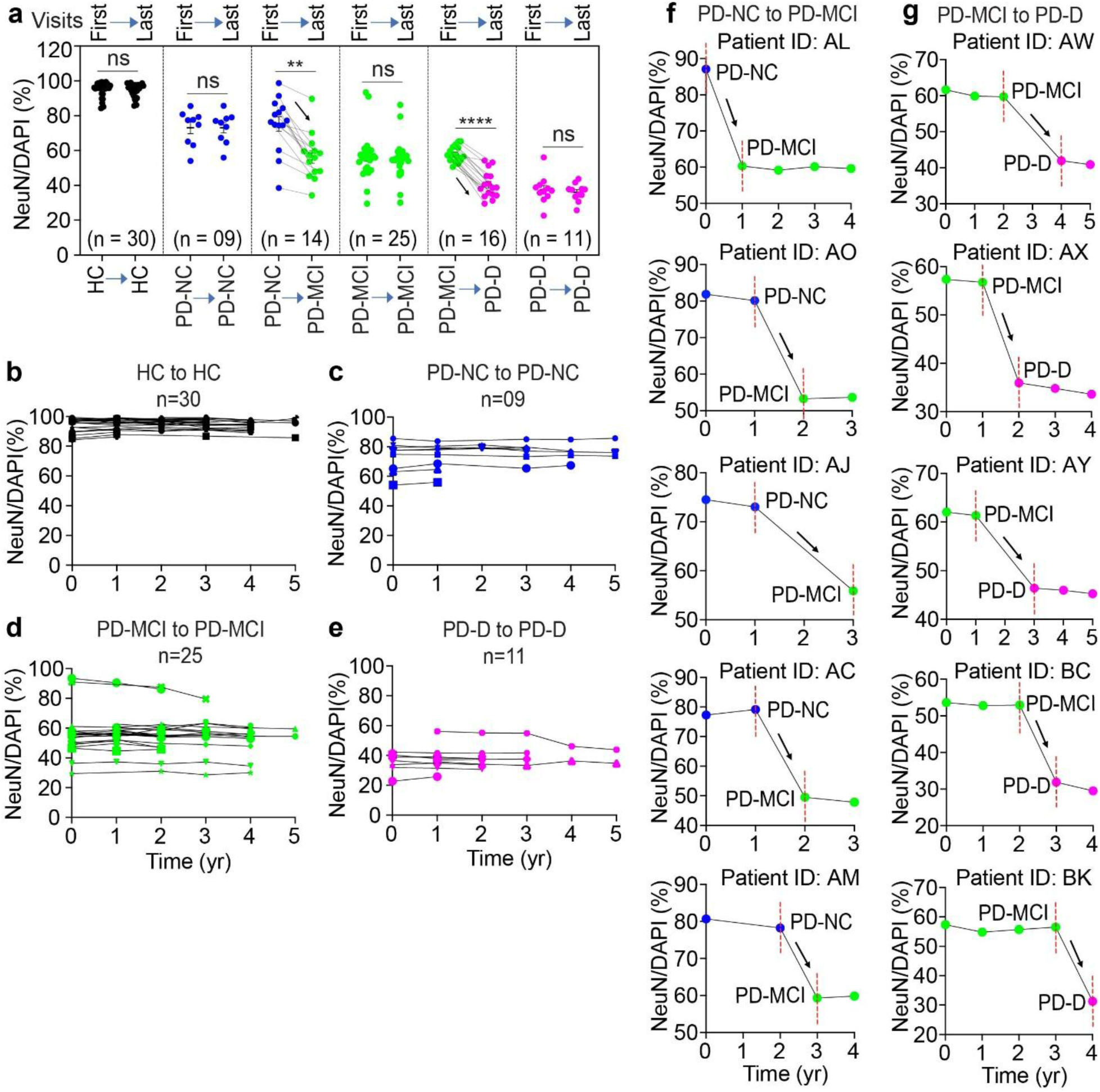
Longitudinal analysis (yearly) for the neurotoxicity of α-syn strains from HC, PD-NC, PD-MCI, and PD-D. Amplified α-syn strains (10 μg/mL) were added to mouse cortical neurons at DIV 7 days. 21 days later neurotoxicity was determined via NeuN counting. (**a**) Neurotoxicity of amplified α-syn strains from longitudinal HC, PD-NC, PD-MCI, and PD-D groups between the first- and last-visit in Cohort I. **(b,c,f)** Yearly mapping of neurotoxicity of amplified α-syn strains from individuals with stable cognitive status. **(d,e)** Yearly mapping of neurotoxicity of amplified α-syn strains from individuals with changed cognitive status. The statistical significance was evaluated via one-way ANOVA with Tukey’s multiple comparisons test. No significant difference (ns) *P* > 0.05, \*\*\*\**P* < 0.0001.

**Extended Data figure 11.**
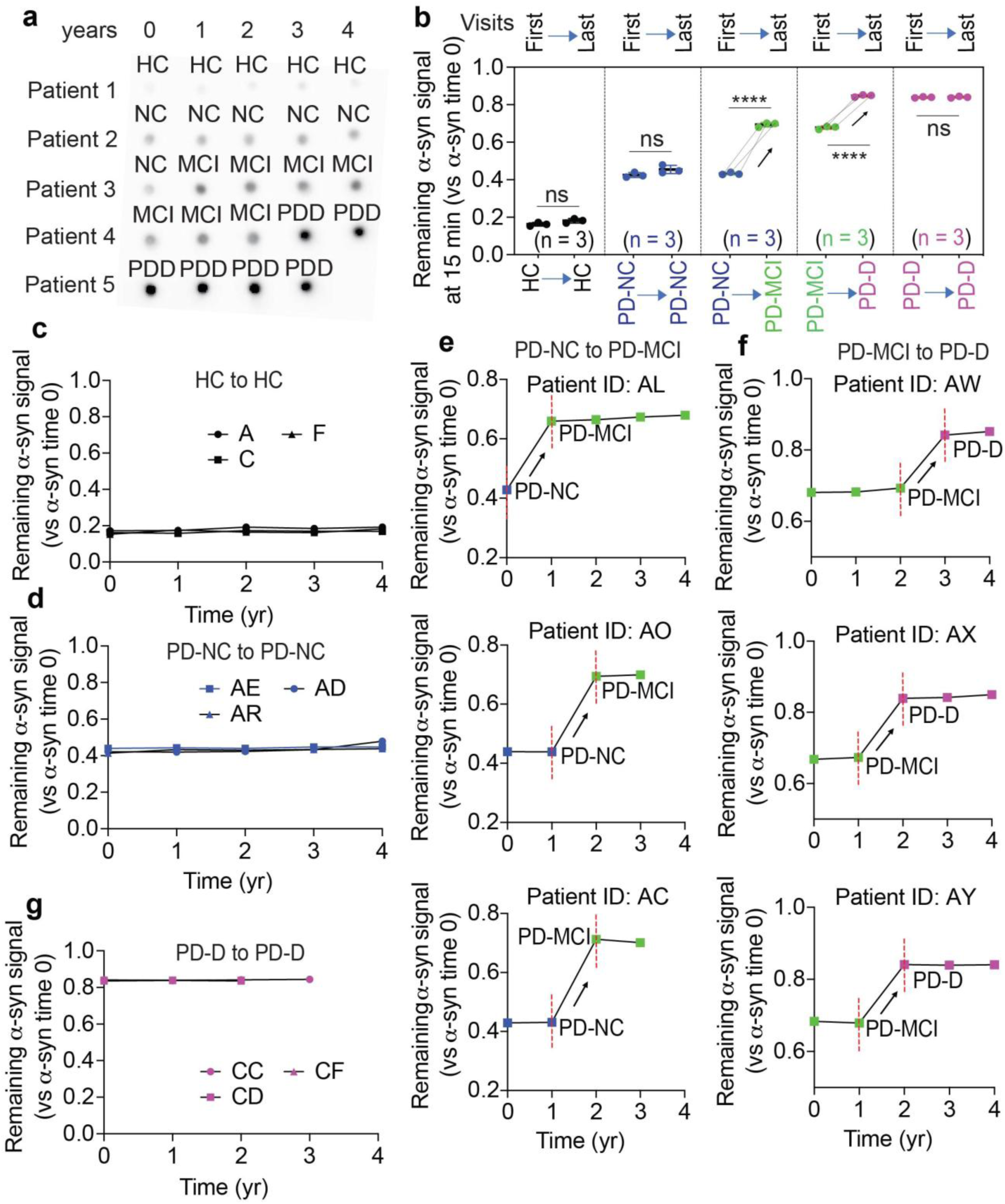
Longitudinal analysis (yearly) of PK digestion of α-syn strains from HC, PD-NC, PD-MCI, and PD-D. (**a**) Dot-blot for PK digested α-syn strains from longitudinal HC, PD-NC, PD-MCI, and PD-D groups between the first- and last-visit in Cohort I. **(c,d,g)** Yearly mapping of the resistance to PK digestion of amplified α-syn strains from individuals with stable cognitive status. **(e,f)** Yearly mapping of the resistance to PK digestion of amplified α-syn strains from individuals with changed cognitive status. The statistical significance was evaluated via one-way ANOVA with Tukey’s multiple comparisons test. No significant difference (ns) *P* > 0.05, \*\*\*\**P* < 0.0001

